# Gut *Clostridia* and *Gammaproteobacteria* metabolize dietary linoleic acid to produce a signature of microbial polyunsaturated fatty acids and associated lipids

**DOI:** 10.1101/2025.09.11.675017

**Authors:** Zongyao Huyan, Nicoletta Pellegrini, Zhen Liu, Ciaran G. Forde, Edoardo Capuano, Josep Rubert

## Abstract

Unlike the well-studied gut fermentation of fibres and proteins, that of dietary lipids remains underexplored. Here, we investigated gut microbial metabolism of linoleic acid (LA), a predominant dietary fatty acid (FA), by the human gut microbiota, integrating isotopically labeled LA (ILLA), isotope tracking lipidomics, metagenomics, and pathway analyses. The findings were further validated within a dynamic gut-simulating environment using the Simulator of Human Intestinal Microbial Ecosystem (SHIME) and secondary analysis of a human dietary intervention trial.

LA exposure led to a broad spectrum of lipids, including LA-derived microbial lipids and others produced as adaptive responses. These included various polyunsaturated FAs (PUFAs) like α-linolenic acid, docosahexaenoic acid, and eicosapentaenoic acid. ILLA tracing indicated that LA was primarily routed to β-oxidation in several species within the class *Clostridia* and *Gammaproteobacteria*, and the intermediates likely facilitated FA synthesis. Temporal patterns revealed that *Clostridia* (e.g., *Enterocloster* spp.) and *Pseudomonas aeruginosa* dominated early-stage LA metabolism, while *Gammaproteobacteria* (e.g., *Klebsiella* spp.) played later roles, producing overlapping lipids that primarily relied on *de novo* FA synthesis. The SHIME confirmed LA bioprocessing over the long term and indicated that a high protein-to-fibre feeding increased PUFA concentration in the colonic condition through its stimulating effects on *Gammaproteobacteria*. The effect of high protein-to-fibre ratio diet on gut microbial PUFA production was confirmed by the results of the human trial.

Overall, this study maps the microbial taxa, pathways, and microbial lipids associated with LA, expanding the knowledge of monoculture experiments and predicting how one of the most abundant FAs is utilized by the human gut microbiota. Our atlas of LA-derived microbial lipids provides a foundation for exploring bioactive microbial lipids that act either locally by exerting their bioactivity on the intestine while concomitantly modulating gut microbes, or systemically, once absorbed, and then released in the bloodstream to target different organs.

## Introduction

A small but significant fraction of dietary lipids, approximately 5–8 g per day, evades absorption in the human small intestine and reaches the colon, where it directly interacts with the commensal microbiota ^1,2^. The nature of this lipid–microbiota interaction depends on the presence of other macronutrients and the composition of fatty acids (FAs) ^34^. Saturated FAs have often correlated negatively with microbial diversity in animal and human studies ^5–7^, whereas diets enriched in unsaturated FAs, especially omega-3 and omega-6 polyunsaturated FAs (PUFAs), tend to foster beneficial gut microbes, such as *Bifidobacterium* and *Lactobacillus* ^6,8,9^. Downstream, these shifts in microbial populations drive metabolic alterations within the gut lumen and, ultimately, affect host metabolic homeostasis ^7,10^.

Beyond altering the structure and function of microbial communities, dietary lipids are also biotransformed into a wide range of microbial lipids that affect local gut physiology ^5,10–14^. Human gut bacteria are known to convert unsaturated FAs into conjugated and hydroxylated derivatives through a process known as biohydrogenation or biotransformation ^11,15^. Both *in vitro* and *in vivo* studies have demonstrated that gut-derived *Bifidobacterium* and *Lactobacillus* metabolize linoleic acid (LA), a predominant dietary omega-6 PUFA in human diets ^16^, into hydroxy and conjugated FAs ^12,17–22^. These microbial lipid metabolites exert local benefits ^11,23^. In mice, conjugated LAs inhibit colorectal tumorigenesis ^11^, while specific hydroxy FAs enhance epithelial barrier integrity via GPR40 signalling ^12^. In a recent study, dihydroceramide phosphoethanolamines, sphingolipids that can be produced by *Bacteroides thetaiotaomicron*, were delivered to host dendritic and epithelial cells, promoting interleukin-10–mediated anti-inflammatory responses via activation of the mevalonate pathway in a murine model ^24^. Despite increasing interest in microbial lipids, the signature of microbial lipids from human gut microbiota during undigested fat metabolism remains uncharacterized.

Based on current understanding of how bacteria metabolize extracellular fatty acids (eFA), gut microbes seem to assimilate dietary FAs into their biomass, thereby modulating interactions with both microbes and the host ^25–27^. Beyond biohydrogenation of PUFAs via enzymes, including myosin-cross-reactive antigen (MCRA) and MCRA-like isomerases/hydratase ^15^, all bacteria can assimilate eFA, and some encode machinery to process them into intracellular metabolic networks ^27^. FA uptake generally occurs by passive diffusion or via dedicated transport proteins. Once in the cytoplasm, eFAs typically bind cofactors like acyl carrier protein (ACP) or coenzyme A (CoA), enabling their entry into elongation and β-oxidation cycles, respectively ^27–31^. These processes yield metabolic intermediates that serve in energy production, lipid synthesis, or other biosynthetic functions ^32–35^. Several bacterial species have been reported to incorporate eFAs into their lipid metabolism, influencing membrane composition, fluidity, and gene expression ^26,33,34,36^. However, it remains unclear which FAs are the preferred substrates of commensal bacteria, what specialized gut microbes are capable of producing microbial lipids, and whether these microbial lipids exert beneficial effects on intestinal cells.

To address some of these knowledge gaps and lay the foundation for others, we designed a study to investigate dietary lipid–gut microbiota interactions within the human gut environment (**Fig. 1A**). We profiled microbial lipids derived from LA by integrating isotope-labeled LA (ILLA) and isotope-tracking lipidomics. Through next-generation shotgun sequencing, we delineated the metabolic reaction networks underlying the microbial metabolism of LA and identified key gut microbes and their functions in LA utilization. In particular, we highlighted the roles of several species within the class *Clostridia* and *Gammaproteobacteria*, as well as their FA metabolic pathways, which resulted in a diverse range of microbial PUFAs (mPUFAs). To better simulate *in vivo* human gut conditions, we employed the Simulator of the Human Intestinal Microbial Ecosystem (SHIME) under feeding conditions differing in fibre-to-protein ratios. This allowed us to validate isotope tracking lipidomics and assess how macronutrient composition and regional colonic environments modulated LA-gut microbiota interactions. In our simulated *in vitro* gut environment, we showed that a protein-rich environment enhanced LA metabolism, specifically through several *Gammaproteobacteria*, resulting in elevated production of LA-derived lipids, particularly mPUFAs, compared to a fibre-rich environment. Lastly, we conducted a secondary analysis of a two-week fully controlled dietary intervention, aiming to confirm that the macronutrient ratio and the composition of fatty acids modulate the abundance of mPUFAs. Overall, this study identified the key microbial contributors, mapped their metabolic functions, and generated a reference atlas of microbial lipids derived from LA metabolism, offering new insights into dietary lipid-gut microbiota interactions.

**Figure 1.**
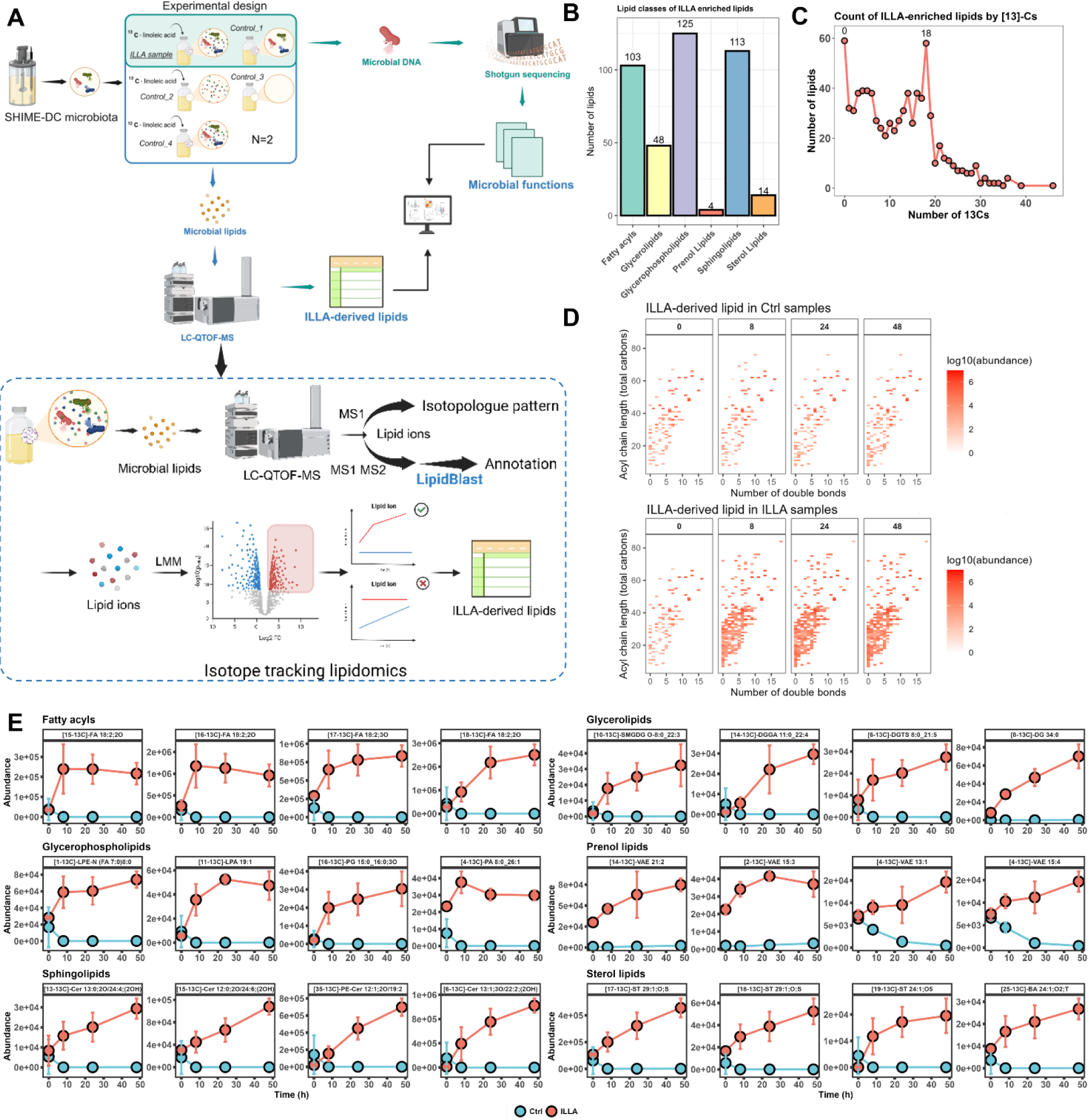
Isotope tracking lipidomics reveals a change in the abundance of ILLA and ILLA-enriched lipids. **A.** The experimental design of ILLA includes ILLA and four control samples, and an overview of the isotope tracking lipidomics workflow for selecting ILLA-derived lipids. The microbial communities (n=2) adapted to the human distal colonic condition in the SHIME were inoculated in the batch culture experiments, and each biological sample was triplicated. The ILLA and control samples contain the following: 1) sterile medium and gut microbiota; 2) sterile fermentation medium and ILLA; 3) sterile medium; 4) sterile fermentation medium, gut microbiota, and non-labeled LA. **B.** The number of ILLA-enriched lipids in different lipid classes. **C.** Number of lipids incorporating isotopically labeled ^13^C atoms as a function of the number of ^13^C atoms incorporated. The two most abundant groups are labeled, including those incorporating 0 and 18 ^13^C atoms. **D.** Double bonds and chain length of ILLA-derived lipids in ILLA and control groups. **E.** Change in time of the abundance of the four most enriched microbial lipids in each lipid class, labeled with the number of ^13^C atoms and annotated lipid names (confidence level II), based on the metric log10(q-value) × coefficient value from the linear model. **Abbreviations:** **ILLA**, isotopically labeled linoleic acid; **Fatty acyls**: **NATau**, N-acyl taurine; **NAPhe**, N-acyl phenylalanine; **NAOrn**, N-acyl ornithine; **NAGlyser**, N-acyl glycine serine; **NAGly**, N-acyl glycine; **NAE**, N-acyl ethanolamines; **FA**, fatty acid; **CAR**, Acylcarnitine; **AAHFA**, acyl acid ester of hydroxyl fatty acid. **Glycerolipids**: **TG**, Triacylglycerol; **SMGDG**, Semino lipid; **MGDG**, Monogalactosyldiacylglycerol; **MG**, Monoacylglycerol; **LDGTS**, Lysodiacylglyceryl trimethylhomoserine; **LDGCC**, Lysodiacylglyceryl-3-O-carboxyhydroxymethylcholine; **DGTS**, Diacylglyceryl trimethylhomoserine; **DGGA**, Diacylglyceryl glucuronide; **DGDG**, Digalactosyldiacylglycerol; **DG**, Diacylglycerol; **ADGGA**, Acyl diacylglyceryl glucuronide. **Glycerophospholipids**: **PS**, Phosphatidylserine; **PI**, Phosphatidylinositol; **PG**, Phosphatidylglycerol; **PE**, Phosphatidylethanolamine; **PC**, Phosphatidylcholine; **PA**, Lysophosphatidic acid; **LPS**, Lysophosphatidylserine; **LPS-N**, *N*-acyl-lysophosphatidylserine; **LPI**, Lysophosphatidylinositol; **LPG**, Lysophosphatidylglycerol; **LPE**, Lysophosphatidylethanolamine; **LPE-N**, N-acyl-lysophosphatidylethanolamine; **LPC**, Lysophosphatidylcholine; **LPA**, Lysophosphatidic acid; **HBMP**, Hemibismonoacylglycerophosphate; **CL**, Cardiolipin. **Prenol lipids**: **VAE**, Vitamin A fatty acid ester. **Sphingolipids**: **SPB**, Sphingoid bases; **SM**, Sphingomyelin; **SL**, Sulfonolipid; **SHexCer**, Sulfatide; **PI-Cer**, Ceramide phosphoinositol; **PE-Cer**, Ceramide phosphoethanolamine; **HexCer**, Hexosylceramide; **GM3**, Ganglioside GM3; **CerP**, Ceramide 1-phosphates; **Cer**, Ceramide. **Sterol lipids**: **ST**, Sterols; **SE**, Sterol esters; **BA**, Bile acids and derivatives; **ASG**, Acylhexosyl. Note: See cross-sectional analyses in **Supplementary Fig. S1B**.

## Results

### The human gut microbiota converts linoleic acid into more unsaturated fatty acids and associated lipids

We collected human stool samples and inoculated them into the SHIME system. After a two-week stabilization period, allowing the microbial community to adapt to the *in vitro* gut environment and having a stable microbial activity, we assessed how the gut microbiota responds to LA using an *in vitro* batch culture. This setup mimics the gut conditions of the human distal colon, characterized by a pH of approximately 6.9, and it ensures that the gut microbiota at taxonomical and functional levels are representative of human stools ^37,38^. Initially, we observed that the gut microbiota rapidly consumed ILLA, with 32% depletion by 8 h and 51% by 24 h. In contrast, only an additional 5% was consumed over the subsequent 24 h (**Supplementary Fig. S1A**). Notably, the abundance of non-labeled LA increased in control but not in ILLA groups.

ILLA exposure resulted in a broad range of lipids produced, as revealed by isotope-tracking lipidomics and linear modelling (**Figs. 1A-B**). To identify the ILLA-derived lipids, we selected ^13^C-labeled annotated lipids that increased in ILLA samples but remained at low abundance or near 0 in control samples across all fermentation time points. All selected lipids are listed in **Supplementary Table S1** and plotted in **Fig. S1**. The most significantly enriched, top 4 of each lipid class based on the metric log10(q-value) × coefficient value from the linear model, are shown in **Fig. 1E**. In total, ILLA increased the abundance of 773 labeled and non-labeled lipids which correspond to 420 specific lipids across 6 major lipid classes: fatty acyls, glycerolipids, glycerophospholipids, prenol lipids, sphingolipids, and sterol lipids, using our criteria (**Table S1**, **Fig. 1B** and **Supplementary Fig. S2C**). These lipids exhibited a wide range of ^13^C incorporations, with non-labeled lipids and lipids containing 18 ^13^C atoms being the most abundant group, mainly attributed to FAs and ceramides (**Fig. 1C** and **Supplementary Fig. S2C**). Furthermore, the produced microbial lipids appeared to raise the number of double bonds in the system (**Fig. 1D**). Notably, 383 out of 420 lipids contained at least 1 ^13^C atom. For instance, 3 of the top 4 most enriched fatty acyls were tentatively identified (level 2) as FA 18:2;2O containing 15, 16, and 18 ^13^C, respectively (**Fig. 1E**). Additionally, 37 of 59 non-labeled lipids were only increased in the form of non-labeled species, i.e., incorporating no ^13^C atoms (**Table S1**). Among all selected lipids, 2 lipids were unambiguously identified using pure analytical standards (α-linolenic acid and ricinoleic acid) ^39^. Additionally, 267 features were annotated at level 2 (accurate mass and MS/MS library spectra) and 151 at level 3 (based on accurate mass and isotopologue pattern) ^39^. The general settings applied to our untargeted lipidomics methods did not provide sufficiently high-quality MS/MS data to annotate monoacyl lipids. However, the MS spectrum provided sufficient information on chain length and double bonds of fatty acids attached to monoacyl lipids, and we decided to report monoacyl lipids from all confidence levels. In contrast, di- and triacyl lipids were only reported at level 2. Lipid names are presented following standard lipidomics nomenclature ^40^.

Taken together, the addition of ILLA to the isolated microbial communities from the SHIME resulted in the production of a broad range of microbial lipids. Notably, this interaction promotes the generation of diverse mPUFAs and increases the overall unsaturation levels. Most lipids, particularly fatty acyls, incorporated either 18 or fewer than 18 ^13^Cs, indicating that their biosynthesis pathways likely involve direct modification of double bonds on the ILLA backbone, acyl chain shortening or elongation, and subsequent incorporation into complex lipids like phospholipids and sphingolipids.

### Linoleic acid exerted time-dependent effects on the gut microbiota, leading to changes in the relative abundance of *Bacillota* and *Pseudomonadota*

The presence of ILLA altered the gut microbial composition and function, as revealed by shotgun metagenomics (**Figs. 2**-**3**). A significant difference in total metagenomic read counts between ILLA and control samples was observed at 24 h (**Supplementary Fig. S3A**). Specifically, total read counts declined in the ILLA group at 24 h, but remained comparable to the control at 8 and 48 h. In the control group, read counts decreased only after 24 h. Microbial diversity also shifted over time. Alpha diversity decreased in both groups, with that of the phylum decreasing at 8 h and species at 24 h (**Fig. 2B** and **Supplementary Fig. S3B**). Beta diversities at the phylum and species levels remained steady in the control group, but they were significantly lower in the ILLA group.

**Figure 2.**
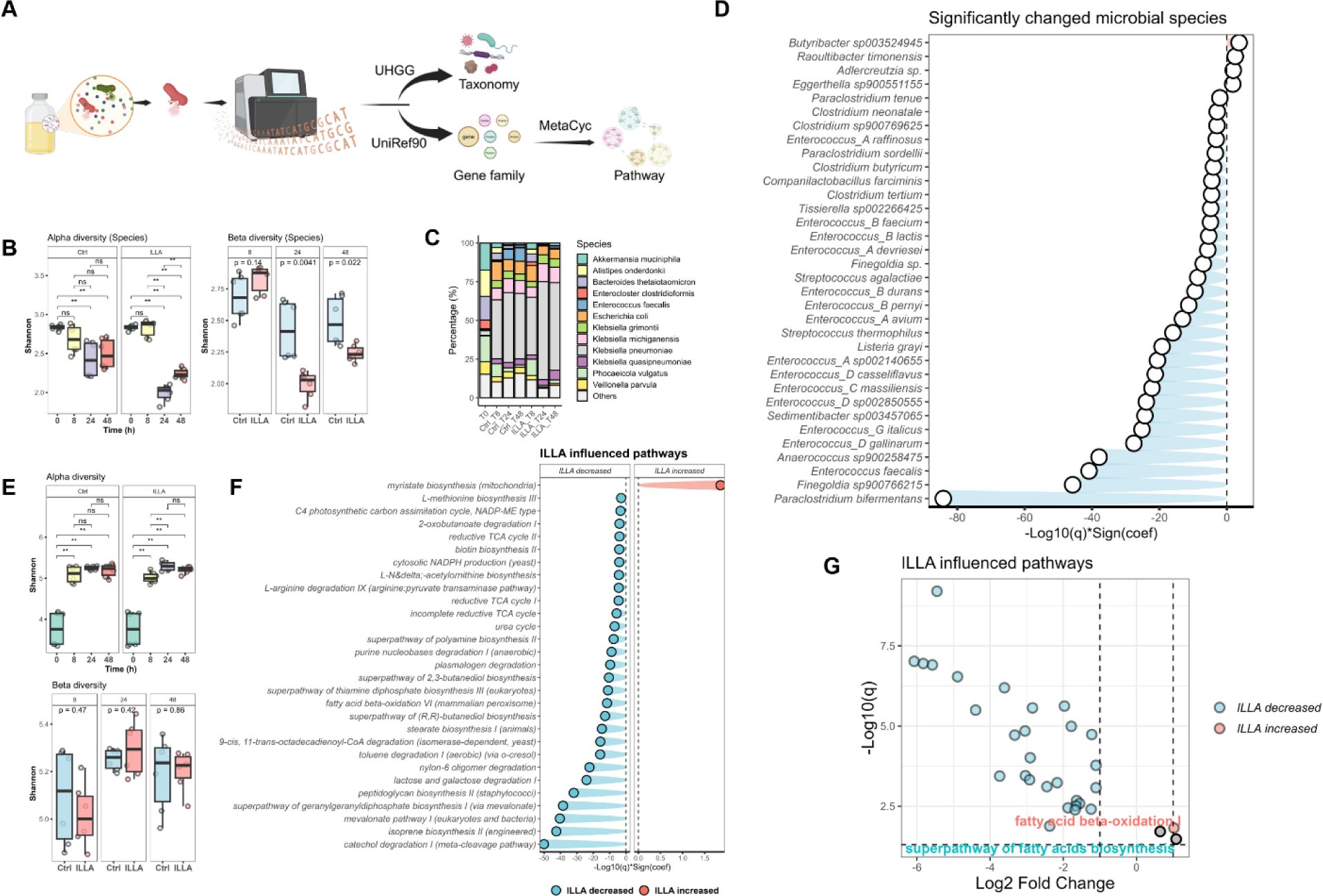
Microbial taxonomic and functional changes reveal the influence of ILLA on the human gut microbiota *in vitro*. **A.** Workflow of metagenomics data processing in the ILLA experiment. **B.** Alpha and beta diversity of gut microbiota at species level in ILLA-treated and control samples (n = 2). Data is expressed as mean ± standard deviation. ns, not significant; *p < 0.05, **p < 0.01, ***p < 0.001, ****p < 0.0001 (One-way ANOVA followed with Tukey post-hoc test). **C.** The most dominant 12 species in the ILLA and control groups (samples containing sterile medium and gut microbiota). **D.** Bar plot illustrating the changes in microbial species in the ILLA and control groups as determined by longitudinal analysis. Statistical significance was assessed using MaAsLin2. Associations were considered significant if they met two criteria: a false discovery rate–adjusted q-value of ≤ 0.05 and an absolute model coefficient greater than 1 (i.e., coefficient > 1 or < –1). **E.** Alpha (**E1**) and beta diversity (**E2**) of gut microbial metabolic potentials in the ILLA and control groups. Data is expressed as mean ± standard deviation. ns, not significant; *p < 0.05, **p < 0.01, ***p < 0.001, ****p < 0.0001 (One-way ANOVA followed with Tukey post-hoc test). **F.** Bar plot showing the significantly influenced metabolic pathways across species, as determined by longitudinal analysis using MaAsLin. **G.** Volcano plot showing the significantly influenced metabolic pathways across species, with fatty acid biosynthesis and β-oxidation pathways labeled, as determined by longitudinal analysis using MaAsLin. Note: see information regarding MaAsLin2 as above.

**Figure 3.**
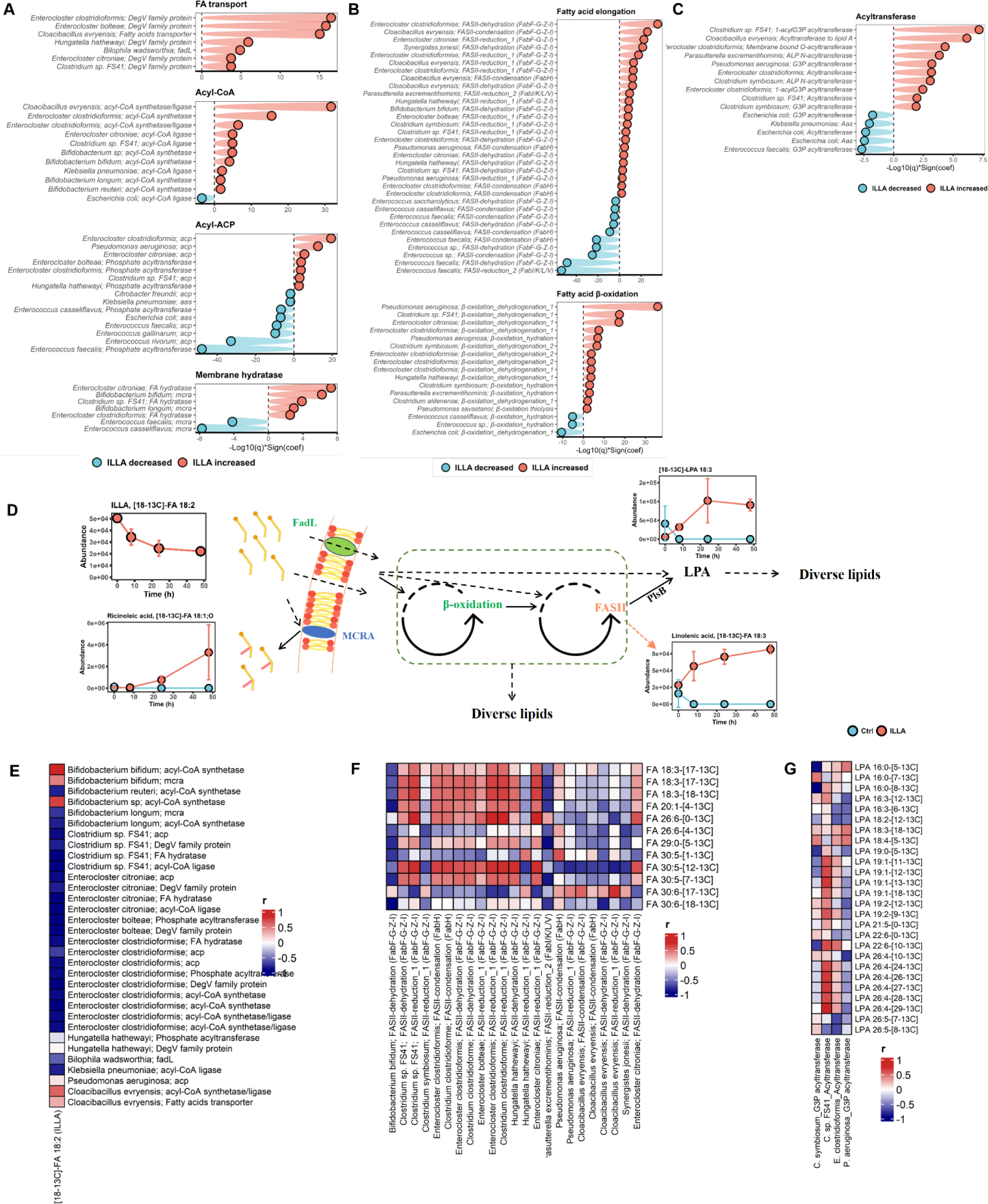
Microbial metabolic potentials and lipidomics data reveal the gut microbes’ role in ILLA metabolism. **A-C.** Bar plot showing the significantly influenced ILLA-enriched gene families encoding proteins involved in fatty acid uptake (**A**), fatty acid elongation/β-oxidation (**B**), and fatty acid transfer (**C**) at 8 hours using MaAsLin. **D.** Hypothesized ILLA metabolic pathways using lipids identified at level 1, such as α-linolenic acid and ricinoleic acid. **E.** Spearman’s correlation of ILLA abundance with gene families encoding proteins related to FA uptake at 8 hours. **F.** Spearman’s correlation of mPUFA abundances with gene families encoding fatty acid elongation at 8 hours. **G.** Spearman’s correlation of ILLA-derived lysophosphatidic acids abundance with gene families encoding glycerol-3-phosphate acyltransferase or potential acyltransferase for glycerol-3-phosphate acylation at 8 hours. Abbreviations: ACP, acyl carrier protein; DG, diacylglycerol; FASII, type II fatty acid synthesis; MCRA, myosin cross-reactive antigen; G3P, glycerol-3-phosphate; GL, glycerolipids; PL, glycerophospholipids; SL, sphingolipids. Note: see information regarding MaAsLin2 in Materials and Methods.

Batch culture protocols utilize a medium rich in proteins, a condition known to favour the proliferation of some species within the class *Gammaproteobacteria*, such as *Escherichia coli* and *Klebsiella* spp. ^41^. Indeed, the microbial composition rapidly shifted toward predominant abundance of *Gammaproteobacteria* members, primarily *Klebsiella* spp., and its belonging phylum *Pseudomonadota* (**Fig. 2C** and **Supplementary Fig. S3C**). The relative abundance of *Pseudomonadota*, mainly driven by *E. coli* and *K. pneumoniae* in this case, increased remarkably from 7.6% to 67.4% in the control samples. In the ILLA samples, *Pseudomonadota* followed a similar trend (60.2% at 8 h) and remained predominant thereafter. In contrast, the initially dominant phylum, *Bacteroidota*, represented by *Alistipes onderdonkii* and *B. thetaiotaomicron*, decreased from 40.6% to 16.6% in control and 18.1% in ILLA samples (**Fig. 2C**). Additionally, other shifts included the emergence of *Enterococcus faecalis* (phylum *Bacillota*) as a newly dominant species (∼8%) in control. Still, it did not appear in ILLA samples after 24 h. Conversely, *Enterocloster clostridioformis* remained a dominant species under ILLA treatment at 8 h (from 5.7 % to 2.3 %). By 24 h, *Gammaproteobacteria* became one of the most abundant in the ILLA group, comprising over 90 % of the total microbial community (**Supplementary Fig. S3C**). Notably, a partial recovery of *A. muciniphila* and *B. thetaiotaomicron* was observed after 48 h post-ILLA exposure.

We next performed both longitudinal and cross-sectional analyses on taxonomic data to identify statistically different microbial taxa modulated by ILLA using Microbiome Multivariable Association with Linear Models (MaAsLin2), as shown in **Fig. 2D** and **Supplementary Figs. S3D-E**. Longitudinally, ILLA decreased the relative abundance of 14 *Enterococcus* and 4 *Clostridium* species, as well as their belonging phylum *Bacillota*. In contrast, it increased the relative abundance of only 4 species, 3 from *Actinomycetota* (*Eggerthella* sp., *Adlercreutzia sp.*, and *Raoultibacter timonensis*) and 1 from *Bacillota* (*Butyribacter* sp.) (**Fig. 2D**). However, a temporary dynamic effect of ILLA was observed in *Bacillota* and its members, increasing in *Bacillota_A* at 8 h followed by subsequent decreases (**Supplementary Fig. S3D2-4**). Similar to trends in diversity, at 8 h, ILLA reduced the relative abundance of 19 species but enriched 78 species (**Supplementary Fig. S3E1**). At 24 h, a greater reduction was observed (109 species decreased their relative abundance, only 3 enriched), while at 48 h, the reducing effect was milder, with 39 species reduced and 52 species enriched (**Supplementary Fig. S3E2-3**).

Overall, these observations demonstrate that ILLA exerts a dynamic and time-dependent effect on the gut microbiota *in vitro*. This effect is characterized by early enrichment of selected species *Pseudomonadota* (e.g., *E. coli*, and *K. pneumoniae*) and subsequent depletion of many Gram-positive (G+) species, such as those within *Bacillota*.

### Linoleic acid elevated FA metabolic potential and modified gut microbial functions mainly in *Clostridia* and *Gammaproteobacteria*

In parallel with the observed taxonomic shifts, we next examined whether ILLA influenced the functional potential of the isolated microbial communities, as reflected by the relative abundance of gene families. Interestingly, despite microbial communities differing between control and ILLA groups, their overall functional capabilities remain conserved (**Fig. 2E**). Notably, beta diversities remained comparable between the two groups across all time points.

To further explore these dynamics, we modelled gene families to species-specific pathways and observed that functional alterations in response to ILLA were limited to a small subset of species. Specifically, out of 230 species modulated by ILLA, only 15 exhibited significant changes in the relative abundance of the pathway (**Supplementary Fig. S4A**). Longitudinal analysis revealed that ILLA primarily enhances the pathway abundance in *Gammaproteobacteria*, particularly in *E. coli, K. michiganensis, and K. oxytoca*, while most pathways in *E. faecalis* increased in control samples (**Supplementary Fig. S4A1**). ILLA triggered more pronounced species-specific pathway increases at 8 h than later (**Supplementary Fig. S4A2-4, S5**). Specifically, some G+ bacteria, *Bifidobacterium bifidum, Bifidobacterium longum, Clostridium aldenense, Clostridium symbiosum, E. clostridioformis,* and *Hungatella hathewayi*, showed elevated cell-wall synthesis (e.g., *UDP-N-acetyl-D-glucosamine biosynthesis* and *peptidoglycan biosynthesis*), energy metabolism (e.g., glycogen metabolism), and amino acid/nucleotide metabolism (e.g., *UMP biosynthesis* and various amino-acid syntheses). In Gram-negative (G-) bacteria, *Pseudomonas aeruginosa* displayed increased phospholipid metabolism (*phospholipid biosynthesis* and *CDP-diacylglycerol biosynthesis*), while *Cloacibacillus evryensis* had higher glycogen and UMP biosynthesis potential. After 8 h, only one, *flavin biosynthesis*, rose in *K. michiganensis* at 48 h in ILLA samples (**Supplementary Figs. S4A4, S5**). Conversely, the relative abundance of most pathways in *E. faecalis* increased in control samples (**Supplementary Figs. S4-5**). Notably, *FA biosynthesis initiation* increased in several *Clostridia,* including *E. clostridioformis* (**Supplementary Fig. S5**).

When gene families were aggregated into metabolic pathways, ILLA increased the relative abundance of only two pathways. ILLA increased the relative abundance of *myristate biosynthesis* in longitudinal analysis and the *reductive TCA cycle I* at 8 h in cross-sectional analysis (**Fig. 2F** and **Supplementary Fig. S4B1**), while markedly reducing isoprenoid biosynthesis (including *isoprene biosynthesis I*, *mevalonate pathway*, and *the superpathway of geranylgeranyldiphosphate biosynthesis*), energy metabolism (*catechol degradation*), and cell wall synthesis (*peptidoglycan biosynthesis*) for the same time point (**Fig. 2F, Supplementary Fig. S4B**). *Myristate biosynthesis* includes FA initiation and elongation, while the *reductive TCA cycle I* processes acetyl-CoA produced from FA β-oxidation. Indeed, we found that both FA biosynthesis (Log2FoldChange = 1.08, q = 0.06) and β-oxidation (Log2FoldChange = 0.65, q = 0.05) increased (**Fig. 2G**).

### *Gammaproteobacteria* and *Clostridia* exhibit metabolic adaptations enabling them to utilize linoleic acid

Given the increase in metabolic potentials associated with LA and the ability of some bacteria to metabolize eFA intracellularly ^27,42^, we next sought to identify microbial species involved in ILLA assimilation and biotransformation. We divided eFA metabolism into sensing, uptake, chain modification, and/or incorporation. The differences in eFA metabolism are primarily determined by species-specific assimilation and chain modification initiation strategies ^27,42,43^. Thus, we explore this process by following the metabolic path and contrasting the mechanisms of G- and G+ bacterial metabolism **(Fig. 3A-D)**, with further insights into known bacterial eFA metabolism discussed later.

In the initial sensing phase, certain bacteria regulate their gene expression through specific sensing systems, like PhoP/PhoQ ^43^. However, ILLA did not increase the relative abundance of any known gene families with FA sensing. Nevertheless, ILLA did increase the relative abundance of gene families encoding functions relevant to eFA metabolism across diverse species (**Figs. 3A-C and Supplementary Fig. S6A, C-F**). Consistently, gene families related to eFA metabolism were prominently elevated at 8 h post-exposure (**Fig. 3A** and **Supplementary Fig. S6A**). ILLA enhanced FA metabolic potentials primarily in *Bifidobacterium* spp., *Clostridia*, and *Gammaproteobacteria*, while in the control group, increased potentials were observed in *Enterococcus* spp. In terms of uptake, G-bacteria typically utilize outer membrane transporters, such as FadL, while G+ bacteria rely on passive diffusion or proteins like DegV and its FA kinase activity ^44^. Here, ILLA increased gene families encoding FadL in G-species (*Cloacibacillus evryensis* and *Bilophila wadsworthia*) and DegV-like transporters in *Clostridia* (**Fig. 3A**). However, these observations were limited to 8 h and were not observed at 24 and 48 h. In addition, the abundance of gene families encoding MCRA and FA hydratase ^15^, membrane proteins reported to produce diverse LA biotransformation, such as ricinoleic acid, increased in *B. bifidum* and *B. longum*, and *Clostridia* (*E. citroniae*, *C.* sp. FS41, and *E. clostridioformis*) in the ILLA group at 8 h exclusively (**Fig. 3A**).

Beyond transport, the relative abundance of gene families involved in acyl-ACP and acyl-CoA synthesis across *Bifidobacterium* spp., *Clostridia*, and some G-species, including *K. pneumoniae* and *P. aeruginosa*, increased in the ILLA group (**Fig. 3A** and **Supplementary Fig S6A, Acyl-CoA**). At 8 h, ILLA increased the abundance of gene families encoding acyl-CoA synthesis in *E. clostridioformis*, *E. citroniae*, *E. bolteae*, *Clostridium* sp. FS41, *H. hathewayi*, *B. bifidum*, *B. reuteri*, and *K. pneumoniae* (**Fig. 3A**), as well as in *P. aeruginosa* (Log2C = 1.33, q = 0.09). Consistent with this, the relative abundance of gene families for CoA synthesis was also elevated in these species (**Supplementary Fig. S6B**). As expected, given the role of acyl-CoA as a substrate for β-oxidation, ILLA also increased gene families encoding FA β-oxidation, specifically initial steps of dehydrogenation and hydration, in these species at 8 h (**Fig. 3B, ‘Fatty acid β-oxidation’**). However, such increases were not observed at 24 h but at 48 h, where *K. oxytoca* and *K. michiganensis* exhibited elevated relative abundance of acyl-CoA synthesis and β-oxidation gene families (**Supplementary Figs. S6A-C**).

In parallel, ILLA increased gene families for acyl-ACP synthesis in three species of *Clostridia* (*E. clostridioformis, E. citroniae,* and *Clostridium sp.* FS41) and *P. aeruginosa* at 8 h, but did not change the relative abundance of these gene families in *E. clostridioformis* and *Klebsiella* spp. at 24 h and 48 h (**Fig. 3A** and **Supplementary Fig. S6A, Acyl-ACP**). Like FA β-oxidation, ILLA increased FA elongation-related gene families in species whose acyl-ACP synthesis gene families were elevated, except for *E. clostridioformis* at 24 and 48 h and *K. pneumoniae* at 48 h (**Supplementary Fig. S6C, ‘Fatty acid elongation’**). Notably, ILLA increased the relative abundance of gene families encoding only the first two steps within elongation in *Clostridia* and the first three steps in *P. aeruginosa* at 8 h, both of which slowed down the last step, the reduction of double bonds at the β position (**Supplementary Fig. S6C1**).

Isotope tracking lipidomics revealed the presence of not only mPUFAs but also complex lipids. We therefore analysed gene families encoding various acyltransferase and lipid-synthesizing enzymes (**Supplementary Fig. S6D-F**). Similarly, the relative abundance of these gene families increased mainly at 8 and 48 h in the ILLA group. Particularly, those associated with the production of phosphatidic acid (PA), LysoPA (LPA), phosphatidylethanolamine (PE), phosphatidylglycerol (PG), phosphatidylserine (PS), and lipid A increased in *E. clostridioformis*, *C. sp.* FS41 and *P. aeruginosa* at 8 h in the ILLA group (**Supplementary Fig. S6D1**). Furthermore, the relative abundance of gene family encoding polyketide synthase, an enzyme recently implicated in microbial sphingolipid biosynthesis ^45^, increased in *E. coli* at 8 h and *K. pneumoniae* at 8 and 48 h in ILLA samples (**Supplementary Fig. S6E**). Additionally, gene families encoding lipid hydrolysis proteins, like ceramidase (*E. clostridioformis*), lysophospholipase (*E. clostridioformis* and *C. evryensi*), and phospholipase C/D (*P. aeruginosa*) increased in the ILLA group. After 8 h, the relative abundance of gene families encoding lipid synthesis increased in only a few *Gammaproteobacteria (E. coli*, *K. pneumoniae*, and *K. michiganensis*) in the ILLA group (**Supplementary Fig. S6F**).

### Differential temporal utilization of linoleic acid by *Clostridia* and *Gammaproteobacteria* and their related microbial lipids

Multiple species exhibited genetic potential for ILLA utilization through various eFA assimilation and FA metabolic pathways. Among them, *Clostridia* and *Gammaproteobacteria* appeared to contribute to ILLA metabolism. While ILLA concentration declined rapidly at 8 h and plateaued after 24 h, the abundance of many ILLA-derived lipids, like mPUFAs or LPAs, continued to increase over time (**Supplementary Fig. S1** and **Supplementary Fig. S7E**). The relative abundance of some *Clostridia,* like *E. clostridioformis,* and their eFA metabolic potentials increased at 8 h (**Fig. 2C**). Subsequently, FA metabolic potentials in some *Gammaproteobacteria*, such as *Klebsiella* spp., became predominant after 24 h. These observations led us to hypothesize that *Clostridia* act as early ILLA utilizers, converting ILLA into diverse acyls, while *Gammaproteobacteria*, like *Klebsiella* spp., participate later, metabolizing both ILLA and available microbial lipids. Since some of these microbes are not commercially available, some are uncultivable bacteria, and single-species cultures cannot replicate the complexity of LA metabolism and the potential cross-feeding interaction in the gut microbiota ^46^, we instead used correlation analysis, incorporating possible functions and microbial lipids to explore the temporal dynamics of ILLA utilization (**Figs. 3E-G**). *In vitro*, ILLA appeared to be metabolized through two pathways: biohydrogenation and endogenous eFA metabolism. The former yielded biohydrogenation products such as [18-^13^C]-ricinoleic acid, while the latter produced [18-^13^C]-linolenic acid, followed by the phospholipid precursor [18-^13^C]-LPA 18:3, a central intermediate for phospholipids incorporating [18-^13^C]-C18:3.

While this correlation analysis may not capture the complete dynamics of the ILLA-microbe interplay, we highlight the significant contributions of *Clostridia* and *Gammaproteobacteria* in the utilization of ILLA. Here, eFA uptake-related gene families in *Clostridia* showed negative correlations with ILLA abundances at 8h. At the same time, those in *Gammaproteobacteria* presented positive correlations throughout, except for the gene family encoding acyl-CoA ligase in *K. pneumoniae* at 8 h (**Fig. 3E** and **Supplementary Fig. S7A**). As suspected, most of the mPUFAs showed positive correlations with FA elongation-related gene families in *Clostridia* at 8 hours and highlighted positive associations with *Gammaproteobacteria* at later time points (**Fig. 3F** and **Supplementary Fig. S7B**). This is further supported by correlations with LPAs, which suggest that FAs produced from FA elongation might be rapidly used (**Supplementary Fig. S7C**). Notably, acyl-ACP in *E. clostridioformis* negatively correlated with ILLA at both 24 and 48 h. However, the long-chain FAs were positively correlated with FA elongation gene families in other *Clostridia* (**Supplementary Figs. S7A1-2**). Furthermore, we also conducted a correlation analysis on the acyltransferase gene families and LPAs (**Fig. 3G**). Gene family encoding the primary LPA synthesis enzyme, G3P acyltransferase, was enriched only in *C. symbiosum* and *P. aeruginosa*. Acyltransferase genes from *Clostridia* correlated with most LPAs, while the acyltransferase in *P. aeruginosa* correlated with a narrower subset. Lastly, to investigate the relationship between specific enzymes and hydroxyl FAs, that is refered as oxidized FAs (OxFAs) here, we correlated gene families encoding potential biohydrogenation membrane proteins, such as MCRA and FA hydratase, and we noticed a positive relationship with [18-^13^C]-C18 OxFAs abundance, which are likely formed via biohydrogenation of ILLA. Here, *mcra* in *B. bifidum* correlated positively with most of [18-^13^C]-C18 OxFAs (**Supplementary Fig. S7D**).

### A protein-rich environment promotes microbial lipid production in the proximal colon by activating fatty acid metabolism in Gammaproteobacteria

The ILLA experiment identified LA-derived microbial lipids and the species and pathways involved in LA biotransformation. However, the static setup, limited substrates, and abrupt environmental shifts may not fully capture human gut microbiome metabolism. To validate the LA metabolism, in a system that addresses the mentioned downsides and avoids interference from the host lipids and absorption, we selected SHIME, a dynamic *in vitro* model that better mimics human colonic conditions than batch fermentations. We controlled exposure to LA and other macronutrients (see more details in the Method section) by feeding the SHIME with media containing varying protein-to-fibre ratios, specifically high-fibre (HF) and high-protein (HP) feeding conditions, both supplemented with equal amounts of fat (**Fig. 4A** and **Table S2**). Sunflower oil, a globally consumed LA source (LA > 50%) ^47^, was utilized as a feeding condition, and from now on, it is referred to as LA supplementation. LA supplementation achieved LA concentrations comparable to those in the ILLA study (∼0.77 vs. 0.8 mg/mL), reflecting the estimated LA amount expected in the human colon (based on approximately 50 g of daily fat intake, of which 10% reaches the colon ^48,49^). The SHIME also mirrored both the proximal colon (PC) and distal colon (DC), hereafter referred to as HF-PC and HF-DC, and as HP-PC and HP-DC, thereby allowing for deep insights into the spatiotemporal interactions.

**Figure 4.**
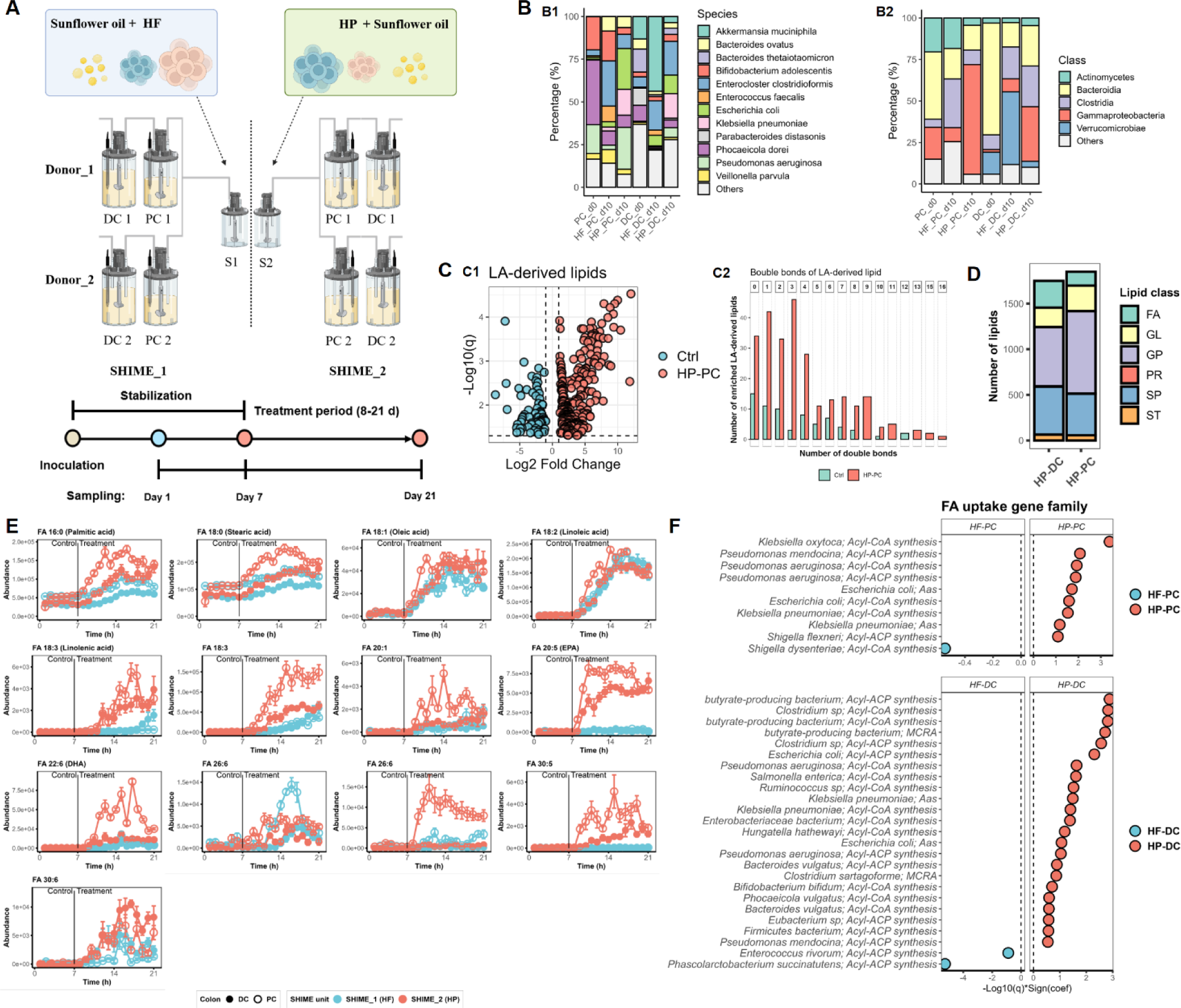
A long-term dynamic fermentation in the TWIN-SHIME confirms the utilization of LA by Gammaproteobacteria. **A.** Experimental design of the TWIN-SHIME system. Four TWIN-SHIME units were inoculated with human gut microbiota (n=2). During a 3-week stabilization period, both units were fed a high-fibre medium (Control). This was followed by two different feeding conditions for 2 weeks: high-fibre/low-protein (HF) or high-protein/low-fibre (HP), supplemented with sunflower oil, making SHIME_1 and SHIME_2. The composition of feeding conditions used in SHIME is shown in **Table S2**. **B.** The most dominant 5 classes (B2), and 12 species (B3) in each SHIME unit at d0 and d10 (c= 2). **C.** (**C1**) Volcano plot showing the significantly changed LA-derived lipids identified in the ILLA experiment in the HP-PC unit before and after sunflower oil supplementation; (C2) distribution of the LA-derived lipids identified in the HP-PC according to their number of double bonds. **D.** Number of lipid species significantly enriched in PC or DC compartments under HP or HF conditions, as selected by MaAsLin. **E.** The abundance of free fatty acids identified using pure standards and LA-derived fatty acids in each SHIME unit against time. SHIME_1 and SHIME_2 represent units fed the HF or HP conditions, respectively. Data is expressed as mean ± standard error of the mean. Two way-ANOVA followed with Tukey post-hoc test was used to compare the differences between each SHIME. The significance is shown in **Table S3**. **F.** Bar plot selecting significantly influenced gene families encoding proteins involved in fatty acid uptake using MaAsLin. Comparison of SHIME units simulating the proximal (PC) or distal colon (DC) under HF versus HP feeding conditions (HF-PC vs. HP-PC and HF-DC vs. HP-DC).

Since the same microbial communities were employed during the ILLA experiment and the SHIME, we could observe that the SHIME basal media modified the initial microbial composition due to a higher presence of fibre in the media (**Fig. 4B**). Despite this, most of the main LA metabolizers, such as *E. clostridioformis* and *K. pneumoniae*, alongside various other *Clostridia* and *Gammaproteobacteria*, increased after feeding with HF and HP supplemented with LA. In addition, most ILLA-derived lipids identified previously, specifically over 90% of the annotated lipids, were also detected in the SHIME experiment using our lipidomics workflow (**Fig. 4E**). A few microbial lipids were already detectable before LA supplementation, indicating that gut microbes may have the capacity to produce them in an LA-free environment. Upon LA supplementation, specific microbial lipids, particularly mPUFAs derived from LA (**Fig. 4C1, Supplementary Fig. S8A,** and **Table S3**). As expected, LA supplementation enabled LA-derived lipids with higher unsaturation degrees (**Fig. 4C2** and **Supplementary Fig. S8B**). Notably, HP-PC enabled the higher abundance of microbial lipids among all SHIME units (**Fig. 4D** and **Supplementary Fig. S8C**).

As in the ILLA experiment, we primarily focused on the presence and abundance of mPUFAs, since they serve as the fundamental building blocks for complex lipids. As expected, most previously identified mPUFAs were not detected before LA supplementation (**Fig. 4E**). After LA supplementation, these mPUFAs started to be released and were present in higher amounts in HP SHIME, particularly in the PC environment (**Table S3**). Notably, upon LA supplementation, we unambiguously identified omega-3 docosahexaenoic acid (DHA) as mPUFA. In the ILLA experiment, the free FA 22:6 was not annotated. However, we identified LPA 22:6 (**Supplementary Fig. S7E**). Additionally, omega-3 eicosapentaenoic acid (EPA), which was not reported as mPUFA in the ILLA experiment, could have been used as an intermediate to be part of complex lipids like PS 8:0_20:5. These mPUFAs, along with many other microbial lipids, were more abundant in HP-PC and HP-DC compared to HF-PC and HF-DC, with HP-PC showing the most significant enrichment (**Fig. 4E**, **Supplementary Fig. S8A,** and **Table S3**). Notably, EPA and DHA were only detected in limited amounts under HF conditions and were not detected before LA supplementation.

The microbial composition confirmed that species, particularly *K. pneumoniae*, *P. aeruginosa*, and *E. clostridioformis,* with eFA metabolic capacity identified in the ILLA experiment, were the most abundant species in HP-PC, which exhibited the highest levels of microbial lipids derived from LA, with similar relative abundances in HP-DC (∼ 40 %). In contrast, *Gammaproteobacteria* accounted for lower than 15 % in the HF-SHIME units, excluding *E. clostridioformis* that remained relatively abundant, 26.4 % in HF-PC and 17.4 % in HF-DC (**Fig. 4B**). In line with compositional data, the relative abundance of gene families encoding eFA metabolism, including uptake, elongation/β-oxidation, and acylation, was elevated in various *Gammaproteobacteria*, including *K. pneumoniae*, *P. aeruginosa*, and *E. coli* in the HP-PC unit (**Fig. 4F** and **Supplementary Fig. S9A-D**). Although similar functional enrichments were also observed in *Clostridia* and *Gammaproteobacteria* in the HF-SHIME units, these occurred at lower abundances compared to HP-PC.

In summary, the dynamic SHIME *in vitro* system provided a more physiologically relevant environment, confirming the observation made in the ILLA experiment. This setup further demonstrated that microbial species, particularly those of *Gammaproteobacteria*, actively utilized LA to produce mPUFAs. The HP feeding condition, particularly in PC, promoted a greater microbial lipid diversity and higher abundances of mPUFAs compared to the HF diet. This was supported by an enrichment in FA metabolic pathways, especially among *E. coli*, *P. aeruginosa*, and *K. pneumoniae*.

### A richer protein diet and a higher proportion of PUFA intake result in a higher abundance of omega-3 PUFAs in stools

Building on our *in vitro* findings, which revealed key mechanistic insights into the metabolism of gut microbes using LA and highlighted the importance of proteins in modulating *Gammaproteobacteria* and thereby stimulating mPUFA production, we aimed to validate these mechanisms *in vivo* through a secondary analysis of a two-week randomized controlled crossover dietary intervention (**Fig. 5A**).

**Figure 5.**
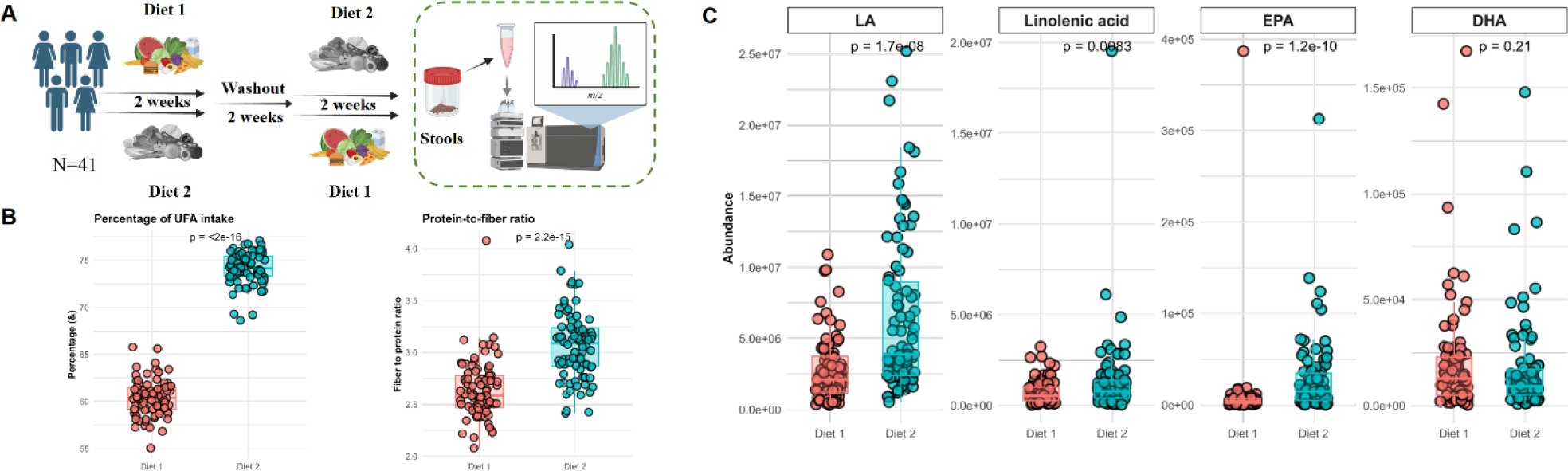
Stool samples collected from a two-week fully controlled intervention (RESTRUCTURE) confirmed the insights obtained from the SHIME and ILLA experiments. **A.** Experimental design of human dietary intervention. Briefly, participants (n=43) were randomly divided into two groups and consumed either diet 1 or diet 2 for 2 weeks. Afterwards, they switched to the other diet for 2 weeks following a 2-week washout period. Stool samples were collected before the intervention on days 5 and 12. Stool lipidomes were determined by the same lipidomic protocol as *in vitro* experiment. **B.** Percentage of unsaturated fatty acids and fibre-to-protein ratio in both designed diets. Significances (p-values) were calculated using the Wilcoxon test, as the Shapiro-Wilk test was also employed, and the variance equality results were reported. **C.** Abundance of linoleic acid, linolenic acid, eicosapentaenoic acid, and docosahexaenoic acid in stool samples collected from participants consuming both diets. Significances (p values) were calculated using Wilcoxon test as Sharpiro-Wilk test and variance equality results suggested.

At the macronutrient level, the two dietary interventions (Diet 1 vs. Diet 2) differed primarily in their protein-to-fibre ratio and relative amount of unsaturated fat content (**Fig. 5B**). Consistent with SHIME results, Diet 2 (higher protein-to-fibre ratio and greater unsaturated fat percentage) led to elevated abundance of linoleic acid in stools (**Fig. 5C**). More importantly, as previously reported, the abundance of linolenic acid and EPA was higher when participants were exposed to high levels of dietary unsaturated fatty acids (p < 0.1, **Fig. 5C**). DHA abundance was comparable between the two groups. Stools, as a byproduct, could not provide us with information about the proximal colon. These observations support the mechanistic connection between undigested proteins and unsaturated fats reaching the colon and stimulating the production of mPUFAs.

## Discussion

Every day, a proportion of dietary lipids escapes digestion and enters the large intestine, where they interact with gut microbiota ^2,50,51^. Due to the overlap of endogenous and microbial lipids, microbial lipid signatures remain largely uncharacterized. Thus, the biological functions of lipids generated by gut microbes are still mostly unknown. To date, most research has focused on microbial biohydrogenation of PUFAs, particularly LA ^50^, yielding hydroxy and conjugated FAs. Beyond this, the metabolic conversion of dietary FAs into diverse lipids and the pathways involved remain elusive. Using LA as a model lipid, in combination with *ex vivo* models, a dietary intervention, isotope-tracked lipidomics, and shotgun metagenomics, we mapped microbial lipid profiles and elucidated the associated pathways.

In short, we identified a wide range of microbial lipids, including mPUFAs, derived from the interaction between the LA and the gut microbiota. Increases in members of the *Bacillota* and *Pseudomonadota* phyla accompanied this enrichment. Within two phyla, *Clostridia* (e.g., *E. clostridioformis*, *E. citroniae*, *E. bolteae*, *C.*sp. FS41, and *H. hathewayi*) and *P. aeruginosa* appear to be the early LA metabolizers processing LA intracellularly, with *Gammaproteobacteria* like *Klebsiella* spp. playing a key role at later stages. Additionally, *Bifidobacterium* spp. utilize PUFAs via membrane-associated MCRA to generate OxFAs, which serve as eFAs for other microbes, facilitating the production of oxidized phospholipids such as OxPEs. Notably, a protein-rich environment stimulates LA metabolism, promoting the formation of mPUFAs and other associated microbial lipids. Consistent with these findings, a secondary analysis of human stool samples revealed that a diet characterized by a higher protein-to-fiber ratio and unsaturated fat promoted a higher abundance of linolenic acid and EPA in stools.

To comprehensively map the production of microbial lipids derived from LA metabolism, we employed ILLA in combination with isotope-tracing lipidomics. Several tools, such as MS-Dial and MetTracer ^52,53^, are available for isotope tracking, relying on the same algorithm. We chose MS-Dial due to its integration with the latest LipidBlast database, which is not included in the MetDHA-MetTracer pipeline. Applying rigorous selection criteria (**Fig. 1A**), we identified 383 lipids across all 6 lipid classes as ILLA-derived microbial lipids (**Fig. S1** and **Table S1**). Interestingly, 59 unlabeled lipids also increased in response to the addition of LA. The majority of ILLA-derived lipids were fatty acyls, glycerophospholipids, and sphingolipids, with those incorporating 18 ^13^C atoms ([18-^13^C]-) being most abundant (**Fig. 1C**). These included mostly FAs and ceramides, with OxFAs constituting the main [18-^13^C]-FAs. We annotated [18-^13^C]-OxFAs incorporating two or more double bonds, such as [18-^13^C]-FA 18:2;2O, which are not products of biohydrogenation of LA but are derived from linolenic acid. Indeed, we also identified [18-^13^C]-linolenic acid as an mPUFA, demonstrating that LA is used as a substrate to generate linolenic acid and its downstream derivatives, such as biohydrogenation products. C18 OxFAs incorporate a broad range of ^13^C atoms, such as [15-^13^C]-FA 18:2;2O and [16-^13^C]-FA 18:2;2O, suggesting formation via chain shortening and elongation, rather than direct biohydrogenation. This chain modification mode is supported by the identification of diverse monoacyl lipids (e.g., [13-^13^C]-NAE 22:5, [10-^13^C]-LPA 22:6, and [6-^13^C]-SPB 16:0;O3) and [18-^13^C]-ceramides, which lack the C18 chain. Additionally, we observed that the LA microbial metabolism generated lipids with higher degrees of unsaturation compared to the LA-free conditions in both short- and long-term *in vitro* experiments (**Fig. 1D**, **Fig. 4C2,** and **Supplementary Fig. S8B**). This shift corresponded with the emergence of linolenic acid, EPA, and DHA (**Fig. 4E**). Although EPA and DHA were not observed in their free form in the ILLA experiment, other complex lipids, such as [21-^13^C]-PS 8:0_20:5 and [0-^13^C]- and [10-^13^C]-LPA 22:6 were identified. Notably, the increased abundance of [0-^13^C]-LPA 22:6 and [0-^13^C]-MGDG 20:5 and 16:2 in ILLA samples suggests mPUFA production as a result of both LA utilization and adaptation ^15,54^ (**Supplementary Fig. S7E**). Subsequently, we consistently observed that a diet with a higher protein-to-fibre ratio and high levels of unsaturated fat consistently stimulated the levels of mPUFAs in both *in vitro* and *in vivo* (**Fig. 4E** and **Fig. 5**). Notably, in our secondary analysis performed on stools from the RESTRUCTURE intervention study, we noticed that mPUFAs were positively associated with the percentage of unsaturated fat intake and a higher dietary protein-to-fibre ratio. In addition, the abundance of other microbial lipids also increased under these conditions, suggesting that undigested proteins and fats are essential for promoting microbial lipid metabolism. Intriguingly, SHIME data indicated the spatiotemporal production of microbial lipids, such as DHA, which was produced in PC, and subsequently, we suspect it was utilized by other gut microbes in DC. This might explain the low DHA levels in our ILLA experiment and *in vivo* secondary analysis (**Fig. 4E** and **Fig. 5C**). Collectively, these results highlight that gut bacteria utilize LA not only through membrane-bound biohydrogenation pathways but also via broader bacterial endogenous FA and lipid metabolic networks.

To identify the gut microbes responsible for converting LA into diverse lipids, we integrated shotgun sequencing and microbial taxonomic and functional profiles were generated. The taxonomic data enabled us to observe the changes in microbial communities, while the latter identified the key species involved in the metabolism of LA. We observed that taxonomic changes in the LA-gut microbiota interaction are temporal. In the ILLA experiment, we used a static batch culture supplied with peptone as the primary carbon source, resulting in a *Gammaproteobacteria*-dominant community centered around *K. pneumoniae*. The addition of LA enriched various *Clostridia*, such as *E. clostridioformis,* at early exposure. In the dynamic *in vitro* model, we observed an enrichment of *Clostridia* in the PC and DC units when LA supplementation was provided for 10 days (**Fig. 4B2**). This enrichment may be related to the cell membrane or wall synthesis, as we observed enhanced functional potentials of cell wall and membrane synthesis pathways in certain *Clostridia* and a *Gammaproteobacteria* species (*P. aeruginosa*) at early LA exposure (**Supplementary Fig. S5**). As LA may exert growth-inhibitory effects on several bacteria ^15^, and stimulate cell wall and membrane molecules, which may better protect the cell integrity and preserve normal functions to co-exist with LA. Because cell wall and membrane molecules are lipid-based or require lipid synthesis, this likely increased demand for lipids, met via *de novo* or incorporating environmental FAs ^27,29,55^. Indeed, we observed an increasing potential of eFA assimilation and FA metabolic function in *Clostridia* (e.g., *Entercloster* spp., *Clostridium* sp., and *H. hathewayi*) and *P. aeruginosa*, using the MaAsLin2 model (**Fig. 3A-C**). As evidenced by other G+ or G-species, certain bacteria can incorporate exogenous FAs into their metabolic networking ^28,56–58^. Our findings suggest a model in which *Clostridia* and *P. aeruginosa* assimilate LA and process it through intrinsic lipid metabolism during the early stages of exposure (8 h). At a later stage, *Gammaproteobacteria*, particularly *Klebsiella* spp., utilize LA following similar pathways, as observed in their enhanced FA metabolic potentials. This progression by *Gammaproteobacteria* is supported by SHIME data, which showed that LA metabolic potential in the later stage (after a 10-day LA exposure) was attributed mainly to *Klebsiella* spp. or other *Gammaproteobacteria* species (**Fig. 4F** and **Supplementary Fig. S9A-D**). These effects were notably pronounced in HP units, which displayed substantially greater abundances of microbial lipids than HF units (**Fig. 4C** and **Supplementary Fig. S8**), likely because proteins favour the expansion of LA-metabolizing *Gammaproteobacteria,* such as *Klebsiella* spp ^41^. Consequently, LA-gut microbiota interactions led to an increase in several *Clostridia* species and a broad spectrum of mPUFAS, including linolenic acid, EPA, and DHA. HP feeding condition in SHIME promoted the expansion of *Gammaproteobacteria,* such as *Klebsiella* spp., further enhancing the LA biotransformation via *Gammaproteobacteria* FA metabolism pathways. While prior studies have shown that some gut microbes can directly incorporate eFAs into cell membrane lipids ^33,34,59^, our isotope tracking lipidomics highlighted that most ILLA-derived lipids contained fewer than 18 ^13^C atoms. This suggests, as summarized in **Fig. 6**, that LA is initially subjected to chain shortening and subsequent elongation before being reused for microbial lipid synthesis.

**Figure 6.**
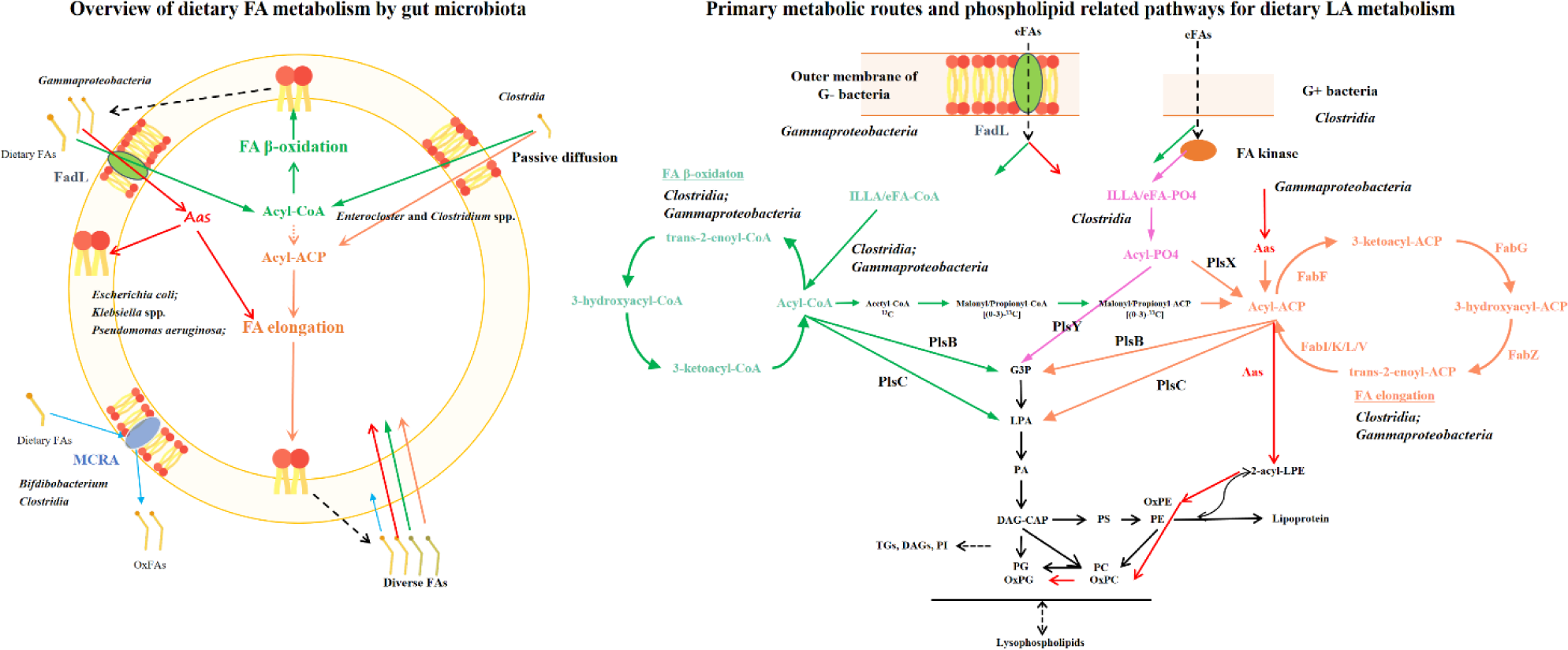
Schematic overview of the microbial metabolism of LA, integrating metabolites and key species identified in this study. The left panel illustrates the overall utilization of dietary fatty acids. In contrast, the right panel displays the metabolic utilization of fatty acids, as well as the production of some of the lipids identified in this study. The proposed gut microbial utilization of dietary fatty acids includes three main steps: **Step 1.** Extracellular fatty acids enter bacteria via different mechanisms: in Gammaproteobacteria, uptake occurs through the membrane transport protein FadL; and, in Clostridia, passive diffusion or DegV protein facilitates fatty acid entry. Once inside the cell, these fatty acids are activated to acyl-CoA. Notably, in Gammaproteobacteria, fatty acids can also be converted to Aas. While conversion to acyl-ACP is a possibility, it has not been documented in either Gammaproteobacteria or Clostridia. In Clostridia, extracellular fatty acids can be converted to acyl-PO4 and then to acyl-ACP, enabling fatty acid elongation. Additionally, enzymes such as MCAR in *Bifidobacterium* and fatty acid hydratase in some *Clostridia* metabolize extracellular fatty acids, releasing modified fatty acids that bacteria can assimilate. **Step 2.** Fatty acids bound to CoA undergo β-oxidation, producing acetyl-CoA. Fatty acids associated with Aas or potentially acyl-ACP enter the fatty acid elongation pathway. The resulting elongated fatty acids are utilized for acylation of glycerol-3-phosphate. Our study indicates that fatty acids derived from the elongation of extracellular fatty acids are more prone to be used for the acylation of glycerol-3-phosphate. Additionally, acetyl-CoA produced from β-oxidation can be converted into malonyl- or propionyl-ACP, serving as substrates for further fatty acid elongation. **Step 3.** The elongated and β-oxidized fatty acids contribute to the biosynthesis of complex lipids, particularly phospholipids: acylation of glycerol-3-phosphate produces lysophosphatidic acid, which can be further acylated to form phosphatidic acid. Oxidized extracellular fatty acids may also be incorporated into phosphoethanolamine, producing oxidized phosphoethanolamine, which can subsequently be converted to oxidized phosphocholine and oxidized phosphoglycerol. The resulting lipids undergo further metabolism within bacterial cells. Abbreviations: **ACP**, acyl carrier protein; **CoA**, co-enzyme A; **eFA**, eFAs; **MCRA**, myosin cross reactive antigen. **PLs**, phospholipids; **SPB**, sphingoid base. **G3P**, glycerol-3-phosphate; **FA**, Fatty acid; **OxFA**, Oxidized fatty acid; **PA**, Phosphatidic acid; **PC**, Phosphatidylcholine; **PE**, Phosphatidylethanolamine; **PS**, Phosphatidylserine; **PG**, phosphatidylglycerol; **DAG**, Diacylglycerol; **DGTS**, Diacylglyceryl trimethylhomoserines; **DGGA**, Diacylglyceryl glucuronides; **TG**, Triacylglycerol; **2-acyl-LPE**, 2-acylglycerophosphoethanolamine; **LPA**, lysophosphatidic acid.

To demonstrate that gut microbes internalize and metabolize LA, we reconstruct step-by-step the uptake of eFA. Initially, *Gammaproteobacteria* activated eFAs via CoA to initiate β-oxidation after uptake through outer membrane FadL or FadL-like proteins ^36,60,61^, as summarized in **Fig. 6**, and in G+ bacteria, such as *Clostridia* ^26,56,62^, ILLA passively diffused across the membrane or was captured by FA-binding proteins, like DegV in *Clostridia* ^62^. Once internalized, our data, which shows elevated abundance of gene families encoding acyl-CoA synthetase/ligase and FA β-oxidation enzymes at 8 h post-ILLA exposure in *Clostridia* (**Fig. 3A-B**), suggests ILLA was bound to CoA. This diverges from monoculture studies, which typically conclude that G+ bacteria do not usually use eFAs for β-oxidation. We propose that both *Clostridia* and *Gammaproteobacteria* employ β-oxidation to process ILLA, generating ^13^C-acetyl-CoA and ^13^C-shorter acyl-CoA, which then serve as substrates for FA elongation. As *Gammaproteobacteria*, evidenced by *E. coli*, lack enzymes to convert acyl-CoA to acyl-ACP ^63, 13^C-shorter acyl-CoA cannot be used for producing new FAs. Therefore, *Gammaproteobacteria* may synthesize mPUFAs via *de novo*, despite an energy-expensive activity ^64^. In *Clostridia*, besides de novo synthesis, there may be an additional pathway that transfers a short acyl group from CoA to ACP for chain elongation. This mechanism was characterized in specific G-species like *Vibrio harveyi* and *Neisseria gonorrhoeae* ^27^. In both cases, FA elongation also requires non-labeled substrates arising from β-oxidation of initially present non-labeled FAs, as most ILLA-derived microbial lipids contain a mix of ^12^C and ^13^C atoms.

The assimilated eFAs also activated the FA elongation pathway. In *Gammaproteobacteria*, upon uptake, eFAs may engage acyl-acyl carrier protein synthetase (Aas), a bifunctional 2-acyl-GPE acyltransferase/acyl-ACP synthase ^28^, thus routing eFA to FA elongation. In *Clostridia*, ILLA or other eFAs may primarily be phosphorylated to acyl-PO_4_ and then converted to acyl-ACP by phosphate acyltransferase (PlsX) that catalyzes the reversible conversion of acyl-PO4 to acyl-ACP ^65^, enabling eFA entry into FA elongation. Notably, in *P. aeruginosa*, we observed an enrichment of gene families encoding phosphate acyltransferase (PlsX, Log2FoldChange = 1.08, q = 0.08), ACP, and FA elongation proteins, suggesting that ILLA taken by *P. aeruginosa*, despite being classified as a *Gammaproteobacteria*, also underwent a pathway via acyl-PO4 and PlsX. These pathways may facilitate the direct elongation of eFAs, such as ILLA. For example, [18-^13^C]-FA 30:6 and [24-^13^C]-LPA 26:4 were likely produced via direct elongation of ILLA using non-labeled ^12^C, possibly by both *Clostridia* and *Gammaproteobacteria*. FA bound to Aas cannot be released until they are acylated to 2-acyl-GPE, indicating that this pathway channels FAs directly into elongation. Nevertheless, we did not detect labeled FA chains shorter than 18 carbons, suggesting that routing eFA into FA elongation is not the predominant pathway. Intriguingly, we observed that the abundance of gene families encoding the last step of FA elongation was abnormally regulated at early exposure of LA **(Fig. 3B**). In this frame, studies also reported that the presence of environmental FA suppresses the gene expression of this step in monoculture ^66,67^. This may explain an increased number of polyunsaturated intermediates starting from early LA exposure (**Fig. 1D** and **Supplementary Fig. S2B**). Here, we identified numerous mPUFAs containing more than 2 double bonds. For instance, we identified α-linolenic acid for the first time as a result of the LA-gut microbiota interactions *in vitro*. Two α-linolenic acid isotopologue species, [17-^13^C]-FA 18:3 and [18-^13^C]-FA 18:3, were identified, and they might be produced via different pathways. As we also identified [18-^13^C]-LPA 18:3, which is produced by acylation of G3P using [18-^13^C]-FA 18:3, [18-^13^C]-FA 18:3 might be produced via FA synthesis. In addition, [18-^13^C]-FA 18:3 might be yielded from desaturation of ILLA via FA desaturase in some *Gammaproteobacteria*, like *P. aeruginosa*, even though only Δ9-FA desaturase, but not Δ15-FA desaturase, has been identified in *Gammaproteobacteria,* like *Pseudomonas putida* ^68^. The abundance of both linolenic acid species rapidly increased at 8 h in the ILLA group (**Supplementary Fig. S7E**), indicating potential contributions from species able to metabolize LA, that is, *Clostridia* and *P. aeruginosa*, to later contributors, *Klebsiella* spp. (**Fig. 3A-B**). We suspected that the identified mPUFAs served as the fundamental building blocks for the synthesis of complex lipids ^29,50^. Among the identified microbial lipids, some, such as [1-^13^C]-FA 30:5, [17/18-^13^C]-FA 30:6, [10-^13^C]-LPA 26:4, and [7-^13^C]-LPA 26:5, peaked at 8 hours and declined thereafter. This suggests that *Clostridia* may take the lead in producing long-chain mPUFAs, which may be crucial to their adaptation to LA. Along with others, these mPUFAs may then be released upon microbial death and recycled. Some bacteria within the order *Lactobacillales* might bypass *de novo* FA synthesis and incorporate eFA into their phospholipids, implying cross-deeding involving FA metabolites from *Clostridia* and *Gammaproteobacteria*. However, as this mechanism has been limitedly studied, we cannot identify the species and lipids within this cross-feeding interaction. While this is underexplored, another cross-feeding route is evident. High relative abundance of functional potential of MCRA in *Bifidobacterium* spp. indicates utilization of newly synthesized unsaturated FAs to produce hydroxy FA, that is OxFAs classified here. OxFAs likely support the synthesis of oxidized phospholipids through Aas-mediated acylation, which acylate 2-acyl-GPE using its bound FAs. For example, [36-^13^C]-PE 18:2_18:1;3O could result from acylation of [18-^13^C]-LPE 18:2 using [18-^13^C]-FA 18:1;O via Aas. PEs were subsequently converted to OxPGs and OxPCs. Due to our conservative criteria, we may miss additional OxFA-containing phospholipids. Importantly, because Aas was found only in *Gammaproteobacteria* in our data, this indicates that oxidized phospholipids came exclusively from them.

As the most abundant identified lipid class, we detected a variety of ILLA-derived sphingolipids. However, the canonical enzyme initiating sphingolipid synthesis, serine palmitoyltransferase, was only detected in *E. coli* and *B. thetaiotaomicron*, without increased abundances in the ILLA group. Interestingly, polyketide synthase has recently been linked to sphingolipid production in *Pseudomonas fluorescens* ^45^, which was enriched in *E. coli* and *K. pneumoniae* in this study. These may underlie the observed ILLA-derived sphingolipids, potentially extending this function to other *Gammaproteobacteria* whose polyketide synthase gene families were also enriched. Additionally, given the primary role of Gammaproteobacteria in the metabolism of eFAs, a diverse spectrum of sphingolipids may be associated with them. Polyketide synthase-like enzymes are also evident in the production of PUFA in marine bacteria and through bioengineering in other bacteria ^69^. Their presence in *Gammaproteobacteria* may contribute to the production of mPUFA. This PUFA metabolic potential may enable the survival and even proliferation of certain *Gammaproteobacteria* in PUFA-rich environments. Further research may consider using knock-out bacteria to gain more insights.

Beyond the reported mechanisms, due to current knowledge and database, we still miss the metabolic insights into many identified ILLA-derived microbial lipids, including fatty acid ester of hydroxyl fatty acids (FAHFAs), fatty amides (FAEs), like N-acyl ethanolamine (NAE), N-acyl glycine (NAGly), and N-acyl tauine (NATau), prenols, and sterols. These lipids incorporate a considerable number of ^13^C atoms, demonstrating their LA origin. In humans, FAEs are likely produced via direct conjugation between FAs and amino acids, catalysis of N-acyltransferase, or degradation of lipids ^70–72^, while their microbial synthesis remains unclear. A similar case happened in FAHFAs, which were first identified around 15 years ago and observed in human intestinal fluids ^73^. Based on the correlation analysis, *Blautia sp.* is considered a producer, but no data have elucidated its FAHFA metabolic potential yet ^73,74^. In our data, we have detected four prenols, all of which are vitamin A fatty acid esters (VAEs). Notably, they are all monoacyl lipids, and one of them [14-13C]-VAE 21:2 likely contains an ILLA-derived FA. To the best of our knowledge, no known biosynthetic route is available. Several sterol lipids, such as [19-^13^C]-ST 24:1;O5 and [24-^13^C]-BA 24:1;O2;T, were identified. These likely arise through the incorporation of ^13^C-acetyl-CoA from β-oxidation, which enters diverse metabolic networks, although no microbial sterol synthesis pathways were elevated in our data. Given the high relative abundance of *Gammaproteobacteria*, they may represent the primary contributors of these sterols. Many of these lipids detected in our study are potentially bioactive and can interact with the human intestine. Bioactive PUFAs like linolenic acid, EPA, and DHA, and in the gastrointestinal tract FAEs are crucial in regulating local physiology and have been associated with immunomodulatory functions ^70,75^, but NAEs were associated with malnutritional status in children^76^. FAHFA has been linked to improving glucose tolerance, anti-inflammatory effects, and anti-cancer potentials, which have been recently reviewed ^77^. However, it is not clear whether their presence in the gut will have local or systemic effects. We also identified lysophosphatidylcholines (LPCs) derived from ILLA and detected them among other LPCs in SHIME. Recent data demonstrates that intestinal microbial-derived LPCs can enter the host lipid circulation and exert beneficial effects on the central nervous system via the gut-brain axis ^14,78,79^. Nevertheless, it remains unclear which microbial LPCs can enact the beneficial effects, although diverse PUFA-containing LPC may benefit the host such as support brain myelin integrity ^80^. Additionally, microbial sphingolipids have both local and systemic health effects ^24,78,79,81^. Some LA-derived microbial lipids may exert health benefits, whereas their presence must be considered within the broader context of the gut ecosystem. For instance, the enrichment of these lipid types co-occurs with the expansion of *Clostridia* and *Gammaproteobacteria*, which are often associated with dysbiosis and inflammatory risk ^82^.

While this study unveils valuable mechanistic insights into how gut microbes metabolize LA, several limitations highlight avenues for future research. One of the key limitations of this study is that the LA metabolism by gut microbiota has not been confirmed in a strict LA-controlled human intervention trail. We were able to perform a secondary analysis on human stools collected for the RESTRUCTURE study, correlating mPUFA production with high UFA and protein intake. During data analysis, we primarily focused on the bacteria’s FA metabolic pathways that are reported to incorporate eFAs, while overlooking eukaryotic microorganisms, such as fungi. Additionally, some complex lipid-containing molecules, such as liposaccharides and lipid A, were not detected, and we also missed potentially other non-lipid metabolites that incorporate ^13^Cs from ILLA. For example, acetyl-CoA, produced from FA β-oxidation, enters diverse metabolic networks, including the citrate cycle. Lastly, in our in vitro study, two donors were selected in alignment with the capacity of the SHIME system and the cost of the scientific materials. Power analysis based on our experimental design (α = 0.05 and β = 0.2) suggests that a sample size of 3-5 donors would be optimal for future *in vitro* studies to enhance statistical robustness.

## Conclusion

Taken together, our findings provide a comprehensive understanding of the LA-gut microbiota interactions, describing the modifications in the chemical space and key microbial contributors. Upon LA exposure, we identified a wide range of microbial lipids that were either directly synthesized from LA or produced as an adaptive response to LA. This extensive metabolic remodelling likely reflects a microbial strategy to cope with LA. To tolerate and metabolize LA, gut bacteria broadly activate FA metabolic pathways. However, the expression of specific genes, particularly those involved in the last step of FA elongation, was likely slowed down, resulting in the accumulation of PUFA intermediates, especially those with more than two double bonds, such as linolenic acids, DHA, and EPA. The key microbial contributors to these processes were species within the class *Clostridia* and *Gammaproteobacteria*. These taxa displayed temporal differentiation in their metabolic activity. *Clostridia*, including *Enterocloster* spp. acted primarily in the early stages of LA exposure, while *Gammaproteobacteria*, such as *Klebsiella* spp., contributed predominantly at later stages, despite producing overlapping mPUFA. We proposed that FAs were likely produced via *de novo* synthesis based on our isotopically labeled evidence. Furthermore, the PC environment, characterized by a pH close to 6, combined with high levels of proteins, favoured microbial lipid production, including various omega-3 mPUFAs. This suggests a region-specific modulation of microbial lipid metabolism in the gut. In addition, fermentative species, such as *Bifidobacterium longum*, may contribute to the further processing of PUFAs through the biohydrogenation mechanism, which involves membrane-associated enzymes like MCRA. Based on this, we infer the pathways of gut microbial bioprocessing of LA (**Fig. 6**). Overall, our study presents a detailed atlas of the LA metabolism by the human gut microbiota, revealing the formation of the mPUFAs, associated microbial lipids, and key microbial players, that is, several *Clostridia* and *Gammaproteobacteria*, involved. These insights into how specific gut microbes utilize extracellular FAs provide novel concepts of bacterial metabolism of FAs and the resulting lipids from this interaction lay a foundation for future research and physiological exploration targeting human gut lipids in health and disease. We expect this new knowledge about mPUFAs to unlock innovative ways to improve metabolic health, attenuate intestinal inflammation, and create personalized nutrition strategies through food-microbiome approaches.

## Acknowledgements

We thank all the dedicated volunteers who participate in this study. We acknowledge our colleague Geert Meijer and Christos Fryganas for help with setting up the liquid-chromatography. The author Zongyao Huyan thank China Scholarship Council for the PhD scholarship (CSC no. 201906300025). This research is supported by the Dutch Top-Consortium for Knowledge and Innovation Agri & Food (TKI-Agri-food) Project Restructure; (TKI 22.150). The ‘RESTRUCTURE project is a public-private partnership on precompetitive research on the influence of food texture and eating rate on energy intake.

## Competing interests

The authors declare no competing interests

## Methods and materials

### Chemicals

ILLA ([18-^13^C]-LA, purity: >95%) and pure analytical standard LA (i.e. [18-^12^C]-LA, purity: >99%) were purchased from Larodan (Solna, Sweden). Arabinogalactan, pectin, xylan, starch, and the adult SHIME growth medium (PD-NM001B) were purchased from ProDigest (Ghent, Belgium). Unless stated otherwise, all other chemicals were of analytical grade and obtained from Sigma-Aldrich (St. Louis, MO, USA).

### Ethics

The Medical Ethical Committee (METC) of East Netherlands declared that the *in vitro* fermentation of this study (ID: 2023-16671) does not fall under the Medical Research Involving Human Subjects Act (WMO), which requires assessment by the METC of the East Netherlands or another recognized medical-ethical review committee. The subjects that provided a fecal sample for SHIME were deemed not to be subject to acts or any conduct that is subject to the Medical Research Involving Human Subjects Act. The human clinical trial was approved by METC of the East Netherlands (Dutch: Oost-Nederland), The Netherlands (NL83462.081.23). Trial was registered at ClinicalTrials.gov, identifier: NCT06113146.

### Gut microbial fermentation of ILLA

#### Adaption of gut microbiota to different colonic regions

To avoid interference and remove initial dietary lipids possible affecting the results, we first adapted the human gut microbiota to SHIME (ProDigest, Belgium) for 3 weeks. Detailed SHIME setup followed our previous study ^83^.

The adaptation started with the inoculation of the SHIME with faecal inoculums from 2 healthy volunteers, who are non-smokers, aged 26 and 40 years old, with no history of antibiotic treatment for at least 6 months. To adapt each microbial community separately to the environmental conditions of colon regions and to remove lipids from the medium, a three-week stabilization was conducted. During this phase, the system was fed the SHIME basal medium (1.2 g/L arabinogalactan, 2.0 g/L pectin, 0.5 g/L xylan, 0.4 g/L glucose, 3.0 g/L yeast extract, 1.0 g/L special peptone, 3.0 g/L mucin, 0.5 g/L l-cysteine-HCl, and 4.0 g/L starch) three times daily. Following the stabilization period, the colonic fermentation was initiated by introducing the prepared inoculum within 8 h.

#### *In vitro* colonic fermentation model

The experiment was designed for the identification of the LA metabolites by applying ILLA. *In vitro* fermentations were executed using inoculums taken from the DC compartment of the SHIME (see previous section) and ILLA.

The *in vitro* colonic fermentation of ILLA was conducted in sterilized penicillin flasks, each containing 6.3 mL autoclaved basal medium (composed of 2.0 g/L NaHCO_3_, 2.0 g/L yeast extract, 2.0 g/L special peptone, 1.0 g/L mucin, 0.5 g/L L-cysteine HCl, and 2.0 mL/L Tween 80). Additionally, the basal medium consisted of 12.34 g/L K_2_HPO_4_ and 10.88 g/L KH_2_PO_4_, with a pH of around 6.8-6.9.

Subsequently, each flask received 0.7 mL of inoculum obtained from DC compartment of the SHIME, resulting in a final LA concentration of 0.85 mg/mL. Two donors were used due to the limited materials. In addition to the experimental treatments (basal medium with inoculums and ILLA added), four groups were prepared as “control” samples using the same procedure: (i) samples containing basal medium, inoculums and LA; (ii) samples containing basal medium with only inoculums (iii) samples containing only the basal medium; (iv) samples containing the basal medium and ILLA. To maintain anaerobic conditions, nitrogen was used to flush the flasks.

The samples were then incubated at 37 °C for 48 h. Whole flasks were retrieved at specific sampling time points (0, 8, 24, and 48 h) and promptly frozen using liquid nitrogen. Each fermentation was run in triplicate. The samples were subsequently stored at -20 °C for further analysis. A pressure meter (Extech SDL700, Extech Instruments, USA) was consistently used to ensure that no leakage occurred from the flasks. It was confirmed that there was no leakage detected in any of the flasks during the assessment.

### Long-term SHIME preparation

A twin-SHIME was designed for the comparison of the influences of diets on the interplay between dietary lipids and gut microbiota. SHIME system is introduced in the text above, and the setup of the SHIME is as follows. Each SHIME unit comprised a combined stomach and small intestine vessel (S1/S2) and was further subdivided into two parallel PC and DC compartments, each inoculated with contributions from the two selected donors. The PC and DC compartments were maintained at volumes of 500 mL and 800 mL, respectively. The entire SHIME system was consistently kept at 37 °C using a warm water circulator (AC200, Thermo Fisher Scientific).

Specifically, two separate vessels were employed to replicate the conditions of the PC (pH 5.6-5.9) and DC (pH 6.6-6.9). To regulate the environment within the PC and DC compartments, the pH was sustained at 5.6−5.9 for PC and 6.6−6.9 for DC. This was achieved by the addition of 0.5 M HCL (acid) or NaOH (base), controlled by the software. The system received three daily feedings with an 8-hour interval. During each feeding cycle, 280 mL of fresh feed (pH 2.0 ± 0.1) was pumped into S1/S2, followed by 120 mL of sterile buffer (12.5 g/L NaHCO_3_). The feeds were then distributed to the series of PC and DC compartments within a 60 min timeframe. The software associated with the system is programmed to maintain a constant pH through automated pH controllers. To ensure anaerobic conditions within the system, a daily nitrogen flush of the headspace (10 min) was incorporated into the programming.

The SHIME experiment was designed to compare two diets, i.e. a HF and HP, following a previous study ^83^. The SHIME setup consisted of two SHIME units, namely SHIME_1 and SHIME_2, each fed with distinct diets, that is, HF for SHIME_1 and HP for SHIME_2, as illustrated in **Table S2** (simulating media). The two diets possess an equivalent quantity of dry matter but differ in their ratios of fibres to proteins. Sunflower oil was used as dietary lipids as it is a predominant LA source ^47^. The amount of sunflower oil used in the experiments was based on the following assumptions calculated based on the dietary fibre consumption: considering an average daily faecal output of 120 g for an individual, a recommended daily intake of 30 g for dietary fibres, a recommended daily intake of 50 g for lipids, and the estimation that 10% of these lipids will enter the colon. Consequently, the ratio between dietary fibres and lipids is approximately 15% ^48,49^.

The SHIME experiment was started by inoculating inoculums from the same 2 healthy volunteers as introduced above. To adapt the microbial community to the environmental conditions of colon regions and to eliminate residual lipids from the inoculums, a two-week stabilization was conducted, as shown in **Fig. 4A**. During this phase, the system was fed the SHIME growth medium (1.2 g/L arabinogalactan, 2.0 g/L pectin, 0.5 g/L xylan, 0.4 g/L glucose, 3.0 g/L yeast extract, 1.0 g/L special peptone, 3.0 g/L mucin, 0.5 g/L l-cysteine-HCl, and 4.0 g/L starch) (PD-NM001B, ProDigest) three times a day. After the stabilization, SHIME were fed with either HF or HP feeding conditions for 2 weeks.

Because the production of short-chain FAs is an indicator of the fermentation processes, the system automatically added a base solution to maintain the pH range of each vessel when it deviated from the specified pH settings. Samples from the fermenter were collected once daily at the same time and stored at -20 °C for subsequent analysis.

### Human intervention

Stool samples were obtained from a crossover human clinical trial (NCT06113146, RESTRUCTURE) conducted by Wageningen University, in which 41 healthy, non-smoking adult volunteers, aged 21-50 years with BMI between 21-27 kg/m^2^, consumed two ultra-processed food diets (**Fig. 5B and Supplementary Fig. S10**). Each dietary intervention lasted 14 days, separated by a 14-day washout period, during which participants resumed their habitual diets.

Participants were randomly assigned to one of two sequences: consuming Diet 1 followed by Diet 2, or vice versa, with 14-day washout in between. All meals were provided ad libitum, matched across arms for energy density, palatability, processing level, and macronutrient content. Food intake (gram and kcal) was recorded at meal, daily and weekly levels. Stool samples were collected on days 5 and 12 of each dietary period, immediately frozen at –80 °C and later processed using the similar lipidomics extraction and analytical workflow we applied to our *in vitro* experiments with modification that indicates in below parts.

### Lipid extraction

Lipids were extracted as previously described ^84^. Briefly, the internal standard (n-heptadecanoic acid) was added to the fermentation samples. Subsequently, the lipids were extracted using a lipid extract solution (1:2, v/v) composed of chloroform/methanol/1.5 % KCl in H_2_O (2:2:1, v/v), according to the procedure of Bligh–Dyer. The lipid phase was obtained, and the solvent was evaporated and then refilled with another solution consisting of isopropanol/methanol/water (65:30:5, v/v/v). Notably, regarding the human stool samples, 200 mg of wet fecal samples were extracted with 0.6 mL methanol and sonicate for 5 min. Lipid phase were extracted after centrifugation at -20 °C for 30 min at 10000 g.

### Untargeted and Isotope-tracking lipidomics of in vitro fermentation samples

Isotope-tracking lipidomics was developed as an extension of an untargeted lipidomics approach, incorporating specialized isotopic tracing techniques during data acquisition, filtering, and interpretation.

For the lipidomics data acquisition, a UHPLC-QTOF-MS system was utilized. The UHPLC system (CTO-40S, Shimadzu, Kyoto, Japan) was equipped with a BEH C18 analytical column (2.1 × 100 mm, 1.7 µm) maintained at 60°C. The mobile phases included: (A) a mixture of water and methanol (95:5, v/v) with 5 mM ammonium formate and 0.1% formic acid, and (B) isopropanol/methanol/water (65:30:5, v/v/v) with 5 mM ammonium formate and 0.1% formic acid. A multi-step elution dual-mode gradient was programmed as follows: starting at 0.0 min (10% B; 0.30 mL/min), the gradient increased to 50% B at 1.0 min (0.30 mL/min), followed by a step to 80% B at 5.0 min (0.30 mL/min). Then, at 11.0 min, it reached 100% B with a slight flow rate increase (100% B; 0.40 mL/min). An isocratic step was maintained for four and a half minutes at 15.5 min (100% B; 0.40 mL/min), and finally, it returned to the initial conditions at 15.1 min (10% B; 0.30 mL/min) and reconditioned up to 17.5 min (10% B; 0.30 mL/min). The sample injection volume was 10 µL, and the autosampler temperature was set at 5 °C.

A QTOF-MS system (LCMS-9030, Shimazu) with an electrospray ionization (ESI) ion source was employed to collect metabolic lipophilic molecules. ESI settings included: nebulizing gas flow at 2 L/min, drying gas flow at 10 L/min, heating gas flow at 10 L/min, and a temperature of 250 °C. The capillary voltage was set at -3.5 kV (negative) or 4.5 kV (positive). The method encompassed a full scan MS range of 50–1200 m/z, followed by the acquisition of product ion spectra ranging from 100–1000 m/z for the twenty most intense ions from the survey spectra throughout the chromatographic run (MS/MS). The collision energy was set at 35 eV with a collision energy spread of ± 15eV. Dynamic background subtraction was enabled. The total cycle time for the MS and MS/MS methods was 0.65 s. Automatic m/z calibration was performed using positive or negative calibration solutions (Agilent, United States). It is important to note that the sequence of the samples was randomized using random number generation. To assess the overall process variability, the lipidomics studies included a set of three technical replicates for each sample and several pooled quality controls obtained by mixing equal volumes of all the samples or each set of samples.

### Untargeted lipidomics of human stool samples

The same BEH C18 analytical column. mobile phase, and multistep programme were used in the analysis of human stool lipidomics, while the UHPLC system (Impact II, Bruker, Bremen, Germany) is different.

A QTOF-MS system (Impact II, Bruker) with an electrospray ionization (ESI) ion source was employed. The negative ionization mode was applied because which can identify more free FAs. ESI settings included: nebulizing gas flow at 2 L/min, drying gas flow at 10 L/min, heating gas flow at 10 L/min, and a temperature of 300 °C. The capillary voltage was set at -3.5 kV negative. The method encompassed a full scan MS range of 50–1200 m/z, followed by the acquisition of product ion spectra ranging from 100–1000 m/z for the nine most intense ions from the survey spectra throughout the chromatographic run (MS/MS). A muti collision energy was set at 20, 35, and 50 eV. Dynamic background subtraction was enabled and set to 0.3 min with a repeat count of 3. The total cycle time for the MS and MS/MS methods was 0.93 s. Automatic m/z calibration was performed using positive or negative calibration solutions (Agilent, United States). It is important to note that the sequence of the samples was randomized using random number generation. To assess the overall process variability, the lipidomics studies included a set of three technical replicates for each sample and several pooled quality controls obtained by mixing equal volumes of all the samples or each set of samples.

### Lipidomics data mining and processing

#### Untargeted and isotope-tracking lipidomics data mining

The data processing was followed as MS-CleanR workflow with modifications ^85,86^. Briefly, an untargeted lipidomics algorithm using retention time (RT, 0.5–16.5 min) and peak picking (100–1200 m/z range) were performed for both positive and negative modes.

All data files were imported to MS-Dial 4.90 using the incorporated lipidomics database (LipidBlast version 68). The MS1 and MS2 tolerances were set to 0.01 and 0.02 Da, respectively, in centroid mode for each. Peaks were aligned on a QC sample as a reference file with an RT tolerance of 0.1 min and a mass tolerance of 0.01 Da. MS-CleanR excluded some molecules with high matching scores as they were identified as fragments rather than individual compounds. For isotope-tracking lipidomics using ^13^C, two distinct QC datasets were employed as reference files. The first QC dataset, derived from samples added non-labeled LA, thereby generating non-labeled molecules, referred to as “parent molecules”. The other QC dataset, originating from samples with ILLA, resulted in ^13^C-labeled molecules, thus referred to as daughter molecules. These parent molecules in the first QC dataset are anticipated to exhibit different daughter molecules in the second QC dataset, characterized by replacing various ^12^C atoms with ^13^C atoms. Other parameters aligned with those used in untargeted lipidomics. To avoid missing ions or isotopes, the data mining was repeated with various parameters, including mass slice, and QC samples that injected during the data collection at various time for over 20 times for each ionization model. Furthermore, the nomenclature for the lipid class follows the definition of the previous publication ^87^.

#### Next-generation sequencing of metagenomics

Microbial genomics DNA samples were extracted from both static and dynamic experiments and sequenced. Briefly, DNA were extracted from samples inoculated with human gut microbiota from two donors with or without ILLA, and SHIME samples from d 7 (d 0 of treatment) and d 17 (d 10 of treatment). The resulting DNA was randomly fragmented to size of 350bp. DNA fragments were end polished, A-tailed, ligated with adapters, size selected, and further PCR enriched. PCR products were purified (AMPure XP system), followed by size distribution by Agilent 2100 Bioanalyzer (Agilent Technologies, CA, USA), and quantification using real-time PCR. Library was then sequenced on NovaSeq 6000 S4 flow cell with PE150 strategy to produce 5 G raw data per sample. Novogene NGS DNA Library Prep Set (Cat No.PT004) is used for library preparation.

#### Bioinformatics analysis of metagenomics

Read quality control and filtering follow previously published protocol ^88^. Several tools, fastqc (v0.12.1), fastp (v0.23.4), and bbmap v39.06, were used to produce quality-filtered data. Taxonomic sequence classification was performed using Kraken2 (v2.0.930) and bracken (v2.9). A species abundance table was generated with Bracken (v2.9) ^89^. To gain more functional insight into the data we have used the HUMAnN 3 pipeline on each sample via the library of UniRef90 and MetaCyc to gain the abundance of gene families and pathways ^90–92^. HUMAnN3 runs the Metaphlan (v4.1.0) program as an intermediate step to assign organism-specific functional profiling, which was performed using the developer-provided MetaPhlAn3 bowtie2 database ^93,94^. Additional scripts embedded within HUMAnN3 were used to align gene family descriptions and merge the 128 original output gene family abundance tables into a single table. Finally, gene families annotated to metabolic reactions were further analysed to reconstruct and quantify metabolic pathways in each sample based on MetaCyc.

#### Statistical analysis

The abundance of lipids was expressed as mean ± standard error of the mean. Statistical comparisons between different treatments were performed using two-way repeated measures analysis of variance (ANOVA) followed by a Tukey post-hoc test with a correction for multiple comparisons using statistical hypothesis testing performed by R (v.4.4.2). Shannon diversities of microbiota and pathways were calculated and expressed as mean ± standard deviation. Statistical comparisons of alpha and beta diversities were performed using t-test and ANOVA, respectively, by R. MaAsLin2 package was used to identify the features with significant changes as illustrated in the text or figure captions, conducted using R ^95^. Lipidomics of samples were analysed with a Bray–Curtis dissimilarity matrix using the vegan package for R. Spearman’s correlation analysis of lipids and lipid metabolism-related pathways and gene families was conducted by Hmisc package for R. The dissimilarity matrix was then projected with non-metric multidimensional scaling to visualize similarities among samples. Permutational multivariate analysis of variance followed by a Benjamini-Hochberg p-value adjustment (vegan package) was furthermore conducted to compare differences among lipidomics from each SHIME unit. A value of p<0.05 was considered statistically significant. The experimental design is made by Biorender@, and the scheme of metabolic pathways of dietary FAs is constructed by Microsoft PowerPoint. All other plots were created by R.

## Notes

### Competing Interest Statement

The authors have declared no competing interest.

